# SOX2-dependent enhancer activation couples epithelial identity to neuronal maintenance

**DOI:** 10.64898/2025.12.06.692710

**Authors:** Sukanya Raman, Akshara Dubey, Anubhav Prakash, Raman Kaushik, Lakshini Kannan, Palak Chugh, Raj K Ladher

## Abstract

The cochlear sensory epithelium and the spiral ganglion neurons it supports arise from a common pool of otic progenitors, yet the neurons remain dependent on the epithelium for guidance and survival long after the two lineages diverge. How that dependency is sustained as the progenitor pool generates increasingly restricted cell types is unknown. Here we show that SOX2 maintains this regulatory continuity. Deletion of *Sox2* from the cochlear epithelium after cochlear neurogenesis does not affect neuroblast generation, delamination and proliferation; however, peripheral axons failed to reach the epithelium and neurons were progressively lost, identifying a non-cell-autonomous requirement for epithelial SOX2. Transcriptomic profiling revealed a SOX2-dependent epithelial programme enriched for neurotrophic and axon-guidance genes, including *Ntf3* and *Bdnf*. In cochlear organoids, acute Sox2 deletion reduced H3K27ac at regulatory elements while H3K4me1 remained stable, consistent with SOX2 maintaining the activity of previously marked enhancers. *Ntf3* and *Bdnf* exemplified distinct regulatory trajectories. At *Ntf3*, SOX2 occupied an enhancer whose activity depended on Sox2, whereas at *Bdnf* it occupied promoter-proximal elements in progenitors before ATOH1 engaged these and a hair-cell-active distal enhancer. Thus, SOX2 links sensory fate specification to epithelial–neuronal communication. More generally, our findings suggest that regulatory competence can persist through lineage restriction, while changing transcription-factor inputs partition its output among descendants to coordinate their development.

## INTRODUCTION

The development of the mammalian cochlea requires precise coordination between two closely connected lineages: the sensory epithelium, which gives rise to hair cells and supporting cells, and the spiral ganglion neurons (SGNs), which provide the afferent connection between the inner ear and the auditory centres of the brain(Appler & Goodrich, 2011). Both populations originate within the early otic epithelium, but their paths separate when otic neuroblasts delaminate and coalesce to form the cochleovestibular ganglion (CVG), from which the cochlear and vestibular ganglia subsequently segregate (Fritzsch et al., 2014; Goodrich, 2016). The cochlear component of the CVG differentiates into SGNs, while cells retained within the otic epithelium establish the prosensory domain and later genaerate hair cells (HCs) and supporting cells (SCs) (Fritzsch et al., 2014; Groves & Fekete, 2012). These lineages therefore begin by physically separating, yet their subsequent maturation depends on their ability to reconnect: differentiating SGNs extend peripheral axons back towards the sensory epithelium, which must supply the guidance and trophic signals required for axon growth and neuronal survival. Axons arrive before the prosensory epithelium differentiates(Bruce et al., 1997; Fariñas et al., 2001; Koundakjian et al., 2007) and remain dependent on it as HCs and SCs emerge, and disruption of this coordination causes defects in cochlear wiring(Fritzsch et al., 1997, 2005; Tessarollo et al., 2004). The epithelial cells producing these signals therefore change identity during the very period in which SGNs depend on them, raising the question of how this neurotrophic programme is sustained across differentiation.

SOX2, a high-mobility group transcription factor, plays an important role in the regulatory hierarchy governing prosensory fate in the cochlea(Kiernan et al., 2005). Its expression begins at the otic placode, persists in the otocyst, and becomes confined to the pro-neurosensory region as the inner ear develops(Dabdoub, 2008; Neves et al., 2011). Genetic studies have established its requirement for neurosensory lineage commitment(Steevens et al., 2017), early neuronal development (Dvorakova et al., 2016, 2020)and hair cell differentiation(Kempfle et al., 2016; Kiernan et al., 2005; Steevens et al., 2019). Within the epithelium, SOX2 regulates fate-defining transcription factors such as ATOH1 and PROX1, with continued expression required for supporting-cell fate but repressed in hair cells (Dabdoub, 2008; Neves et al., 2012; Pan et al., 2013). Notably, conditional deletion of *Sox2* produces a biphasic loss of inner ear neurons, in which neurons initially form and extend projections but subsequently undergo apoptosis(Dvorakova et al., 2016). In that model, however, *Sox2* was deleted in both the epithelium and the delaminating neurons, and the neuronal death was interpreted as a secondary consequence of absent hair cell targets. It was not possible to distinguish a neuronal requirement for SOX2 from a loss of epithelial support, and neuronal death was attributed to the absence of hair-cell targets. It therefore remains unknown whether epithelial SOX2 directly controls the extracellular and neurotrophic signals required for SGN innervation and survival, independently of its role in sensory epithelial differentiation.

Among the neurotrophins best characterised in cochlear development are Neurotrophin-3 (NTF3) and Brain-derived Neurotrophic Factor (BDNF), which promote SGN survival and peripheral axon growth through TrkC and TrkB(Ernfors et al., 1995; Fariñas et al., 2001; Pirvola et al., 1992; Schimmang et al., 2003; Stenqvist et al., 2005). As the prosensory epithelium differentiates their expression is partitioned: *Ntf3* is expressed by SCs, whereas *Bdnf* is expressed mainly by HCs(Fariñas et al., 2001). Loss of either produces stereotyped, region-specific disruptions in SGN survival and afferent wiring (Fariñas et al., 2001; Fritzsch et al., 1997). How expression of *Ntf3* and *Bdnf* is preserved through the transition from the prosensory state to these distinct differentiated cell types, such that trophic support is sustained without interruption, is unknown.

One possible basis for this regulatory continuity could be the enhancer landscape that is established in the progenitor and then reinterpreted by the descendent cell types after differentiation. Enhancers can become competent before their target genes are expressed and remain available as cells become lineage restricted, allowing lineage-specific transcription factors to activate, maintain or silence elements inherited from a common progenitor (Aksoy et al., 2013; Bergsland et al., 2011; Lilja et al., 2017).These regulatory states are reflected, by their histone modifications: H3K4me1 is commonly associated with primed or competent enhancers, whereas H3K27ac is associated with enhancer activity. Comparing how these marks respond to transcription-factor loss can therefore distinguish a requirement for maintaining enhancer activation from a requirement for enhancer competence. SOX2 can engage nucleosomal DNA and has been characterised as a pioneer factor in biochemical and reprogramming studies(Dodonova et al., 2020). Whether SOX2 establishes enhancer competence in the cochlear epithelium or acts on pre-marked elements to maintain their activity, however, remains unknown. We therefore asked whether SOX2 provides regulatory continuity during lineage restriction by establishing the epithelial enhancer landscape or by maintaining the activity of elements carried forward from an earlier prosensory state.

To address this, we conditionally deleted Sox2 from the cochlear epithelium. Loss of epithelial SOX2 disrupted SGN peripheral axon projections and caused progressive neuronal death, despite the initial generation and delamination of neuroblasts. Transcriptomic profiling identified a SOX2-dependent epithelial programme enriched for neurotrophic and axon-guidance genes, including *Ntf3* and *Bdnf*. Chromatin profiling in cochlear organoids showed that *Sox2* deletion altered the enhancer activation mark H3K27ac while largely preserving the underlying H3K4me1 landscape. At *Ntf3*, SOX2 maintained enhancers active in prosensory and SC states. By contrast, *Bdnf* regulatory elements remained primed but inactive in supporting cells and were activated in prosensory region and in HCs. Together, these findings identify SOX2 as a regulator of epithelial-to-neuronal communication and support a model in which enhancer competence is carried forward during lineage restriction, while its transcriptional output is partitioned among epithelial descendants to sustain the development of related neurons.

## RESULTS

### *Sox2* expression in cochlear progenitors is required for spiral ganglion innervation

To specifically dissect the role of SOX2 in cochlear progenitors, we used *Emx2-Cre* knock-in mice crossed with *Sox2*-floxed mice and, for initial characterisation of the recombined domain, a Cre-dependent Rosa26-tdTomato reporter. Emx2-Cre activity begins in the developing inner ear at approximately E11.5 and overlaps the SOX2-positive prosensory domain by E12 (Ono et al., 2014)(Fig. 1A).At E12, control embryos (*Sox2^wt/fl^;R26^tdTomato/+^;Emx2^Cre/+^*) expressed SOX2 normally. Cre activity was not detected in neuroblasts at this stage. *Sox2^fl/fl^;R26^tdTomato/+^;Emx2^Cre/+^* embryos showed a loss of SOX2 from the recombined prosensory epithelium (Fig. 1 *B*). For subsequent analyses, we compared reporter-free littermates, Sox2^fl/fl^; Emx2^Cre/+^ embryos (hereafter Sox2 cKO) with Sox2^wt/fl^; Emx2^Cre/+^ controls. By E14.5, SOX2 was undetectable in the prosensory domain and lateral Kölliker’s organ of the Sox2 cKO (SI Appendix S1 A and B). Marker analysis revealed that at E14.5, Sox2 cKO cochlea show reduced p27KIP1 expression across the apical, middle and basal prosensory domains (SI Appendix S1 A and B). By E15.5, mutant cochleae lacked the organised β-spectrin-positive hair cells and PROX1-positive supporting cells observed in controls (SI Appendix S1 C). Alongside the loss of SOX2 in the cochlear epithelium, sagittal sections of the Sox2 cKO, compared to the controls, showed loss of peripheral axons while NeuN-positive neurons and central axon projections, marked by TUJ1 nevertheless remained evident in the mutant ganglion (Fig1 C&D).

**Fig. 1.**
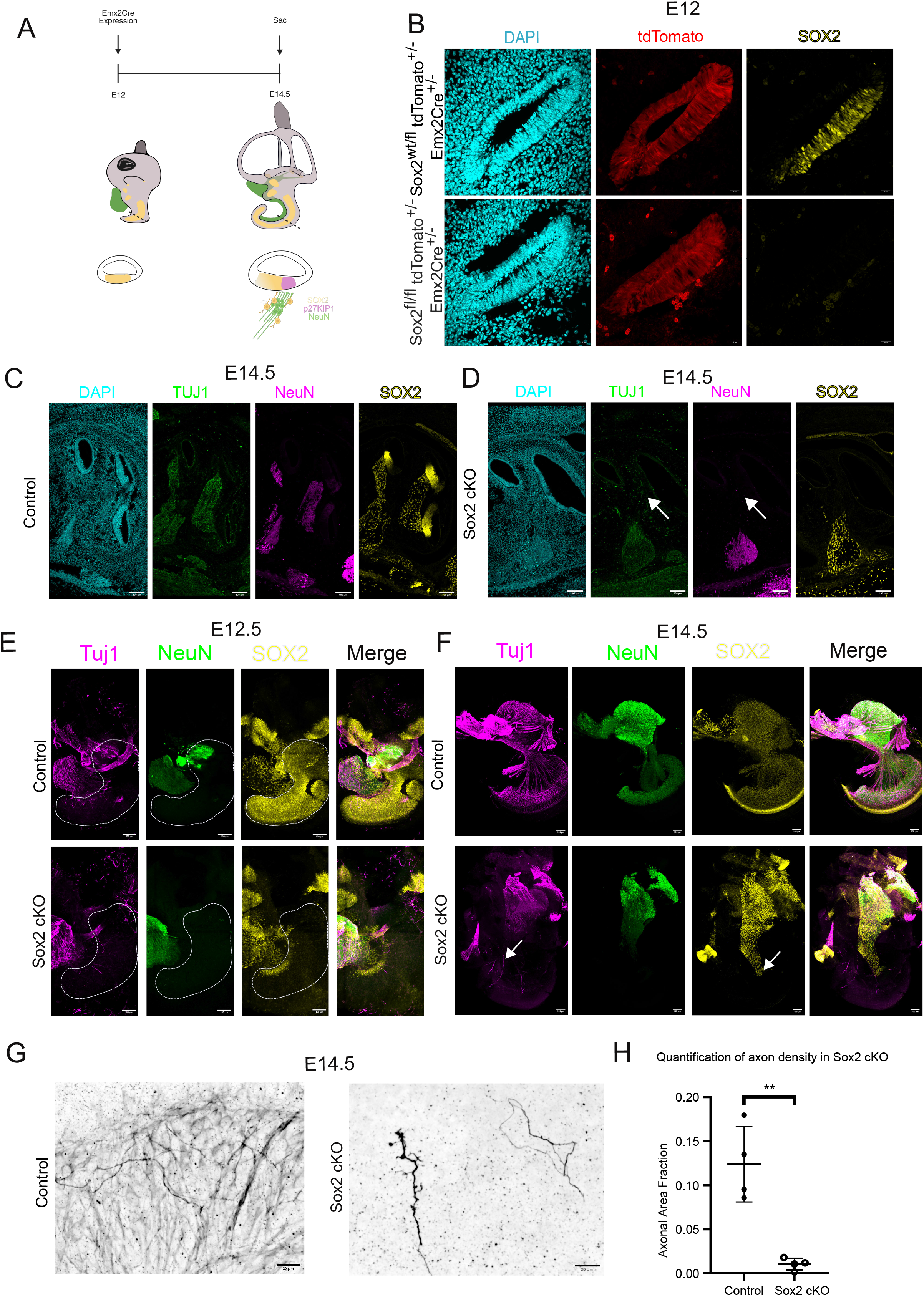
Sox2 deletion from the cochlear prosensory domain disrupts peripheral innervation by spiral ganglion neurons. (A) Scheme of the experimental design. Emx2-Cre activity in the inner ear begins at ∼E11.5 and overlaps the SOX2-positive prosensory domain by E12; embryos were collected at E12, E12.5 and E14.5. (B) Transverse sections through the E12 cochlear duct of control (Sox2^wt/fl^; R26^tdTomato/+^; Emx2^Cre/+^, Upper) and Sox2 cKO (Sox2^fl/fl^; R26^tdTomato/+^; Emx2^Cre/+^, Lower) embryos, labelled for DAPI, tdTomato and SOX2. tdTomato reports the recombined domain; SOX2 is lost from the recombined prosensory epithelium in the cKO. (Scale bar 20 µm) n = 4 embryos per genotype. (C and D) Sagittal sections of the E14.5 inner ear from control (C) and Sox2 cKO (D), labelled for DAPI, TUJ1, NeuN and SOX2. TUJ1-positive peripheral projections toward the cochlear epithelium are present in the control and absent in the cKO (arrows), whereas NeuN-positive nuclei and TUJ1-positive central projections remain in the mutant ganglion. (Scale bar 100 µm.) n = 4 embryos per genotype. (E) Whole-mount E12.5 cochlea with the spiral ganglion intact, control (Upper) and Sox2 cKO (Lower), labelled for TUJ1, NeuN and SOX2. Dashed lines outline the cochlear epithelium. (Scale bar 100 µm.) n = 5 per genotype. (F) Whole-mount E14.5 cochleae, control (Upper) and Sox2 cKO (Lower), labelled as in E. Arrows indicate loss of peripheral axons near the sensory epithelium. (Scale bars, 100µm.) n = 4 per genotype. (G) High-magnification views of TUJ1 labelling (inverted grayscale) of peripheral neurites in the basal turn at E14.5, representative of the fields quantified in H. (Scale bar 20 µm.) (H) Axonal area fraction in control and Sox2 cKO cochleae at E14.5. Each point represents one cochlea; bars show mean ± SD. n = 4 control and 5 cKO. Unpaired t test; **P = 0.002.

We next asked whether the loss of cochlear epithelial identity was accompanied by an early defect in peripheral innervation which starts at E12.5 in the mouse cochlea(Koundakjian et al., 2007). We used whole mount cochlear tissue with the spiral ganglion intact to study innervation across the entire cochlea. At E12.5, TUJ1-positive peripheral neurites extending towards the cochlear epithelium appeared sparser in Sox2 cKO embryos than in controls (Fig. 1E). By E14.5, control SGNs had formed extensive peripheral projections towards the cochlear duct, whereas these projections were largely absent from Sox2 cKO cochlea (Fig. 1 F). Quantification confirmed a marked reduction in peripheral axonal area in Sox2 cKO cochlea relative to controls (Fig. 1G, H).

At E14.5, paint-filled Sox2 cKO inner ears retained the major structures of the membranous labyrinth (SI Appendix S1 D and E), and cochlear duct length did not differ significantly from controls. However, by E16.5, the mutant cochlear duct was significantly shorter (SI Appendix S1 F and G). Thus, the innervation phenotype was detectable before the onset of significant cochlear shortening and in the presence of differentiated neurons and central projections. We therefore next examined whether the loss of peripheral innervation reflected defective neuroblast production, impaired axon extension or a subsequent failure of neuronal survival.

### Sox2 sustains an epithelial-specific programme for neuronal support

Deletion of *Sox2* at E12.5 has been reported to cause only limited neuronal defects (Steevens et al., 2017). However, using *Emx2*-Cre, we found a pronounced loss of peripheral axon projections toward the cochlear epithelium. We thus asked whether this reflected defective neuroblast production, or a failure after neurons had been generated. We first used the Rosa26-tdTomato reporter to trace cells derived from the Emx2 lineage. At E13.5, we detected tdTomato-positive cells, co-labelled with the neuroblast marker, NeuroD1, within the spiral ganglion in both control and Sox2 cKO embryos (2A). The number of tdTomato-positive cells within the ganglion did not differ significantly between genotypes (Fig. 2B). This suggests that there is no defect in neuroblast delamination at this stage.

**Fig. 2.**
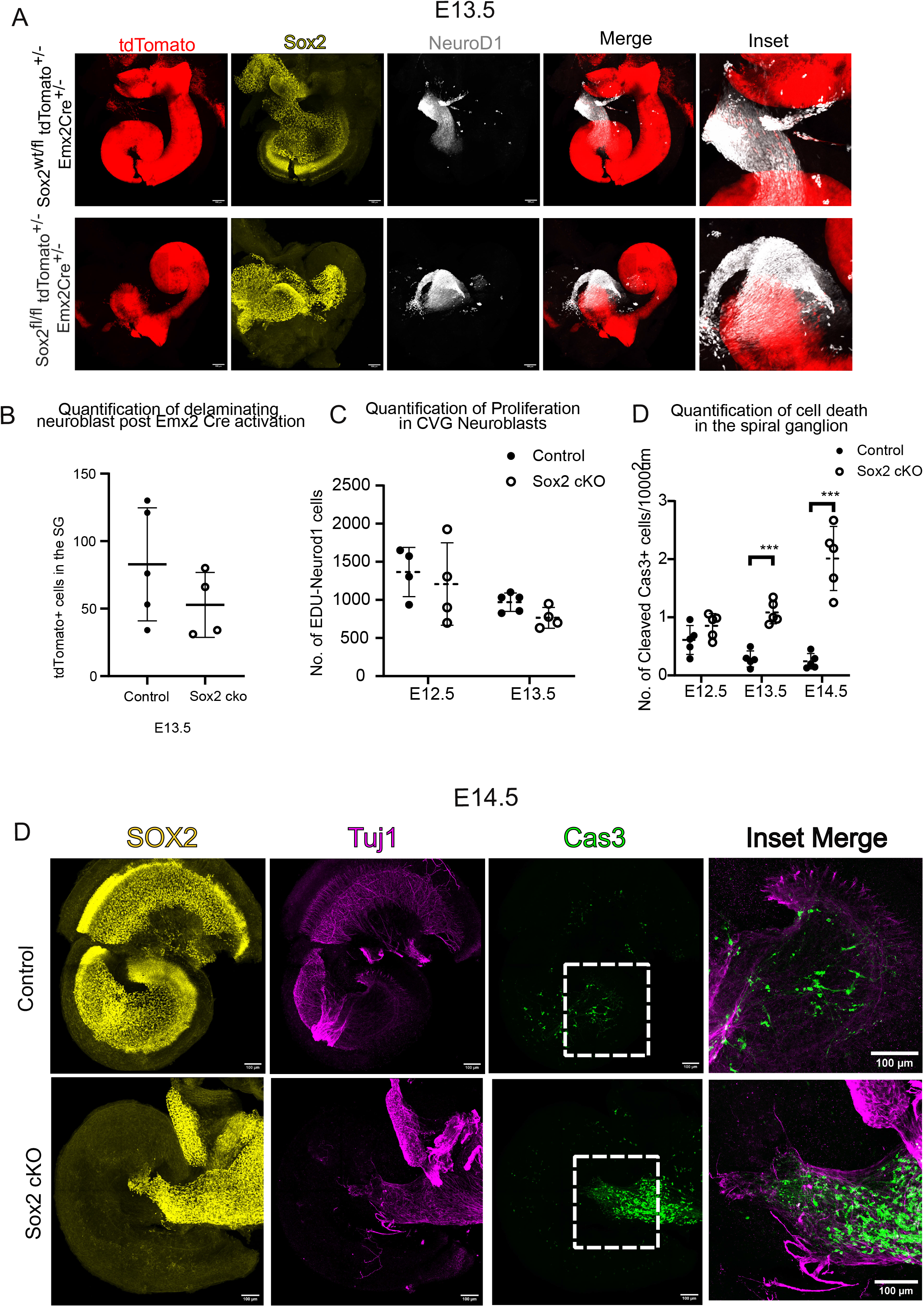
Spiral ganglion neuroblasts are generated and delaminate normally in *Sox2* cKO embryos but are subsequently lost. (A) Whole-mount E13.5 cochleae from control (*Sox2*^wt/fl^; *R26*^tdTomato/+^; *Emx2*^Cre/+^, *Upper*) and *Sox2* cKO (*Sox2*^fl/fl^; *R26*^tdTomato/+^; *Emx2*^Cre/+^, *Lower*) embryos, labelled for tdTomato, SOX2 and NeuroD1. tdTomato marks *Emx2* lineage-derived cells; NeuroD1 marks neuroblasts. *Insets* show the spiral ganglion at higher magnification. (Scale bar 100 µm.) (B) Number of tdTomato-positive, NeuroD1-positive cells in the spiral ganglion at E13.5. Each point represents one embryo; bars show mean ± SD. *n* = 5 control and 4 cKO. Unpaired t test; *P* = 0.24. (C) Number of EdU-positive, NeuroD1-positive cells in the cochleovestibular ganglion at E12.5 and E13.5, quantified from serial sections (*SI Appendix*, Fig. S2*A*). EdU was administered 6 h before collection. Each point represents one cochlea; bars show mean ± SD. *n* = 4 per genotype per stage.; *P* = 0.63 at E12.5 and *P* = 0.0561 at E13.5. (D) Density of cleaved-caspase-3-positive cells in the spiral ganglion at E12.5, E13.5 and E14.5, expressed per 1,000 µm². Each point represents one embryo; bars show mean ± SD. *n* = 4 per genotype per stage. Unpaired t test; \*\*\**P* = 0.0001; the difference at E12.5 was not significant (*P* = 0.13). (E) Whole-mount E14.5 cochleae with the spiral ganglion intact, control (*Upper*) and *Sox2* cKO (*Lower*), labelled for SOX2, TUJ1 and cleaved caspase-3. *Insets* show merged TUJ1 and cleaved caspase-3 signal in the ganglion. (Scale bars, 100 µm; *Insets*, 100 µm.)

Spiral ganglion neuroblasts are generated between approximately E9.5 and E12.5, after which the major phase of neurogenesis is complete (Koundakjian et al., 2007; Wikström & Anniko, 1987). As Emx2-Cre-mediated recombination overlaps the final period of otic neurogenesis, we asked if neuroblast proliferation was affected in Sox2 cKO. Serial sections through the cochleovestibular ganglion were labelled for EdU and NeuroD1, and EdU-positive, NeuroD1-positive cells were quantified at E12.5 and E13.5 (Fig. 2C; SI Appendix S2). Their numbers did not differ significantly between control and Sox2 cKO embryos at either stage. Thus, at the stages examined, Sox2 deletion did not produce a detectable reduction in neuroblast proliferation.

We next asked whether neurons that had formed subsequently failed to survive. At E12.5, the density of cleaved-caspase-3-positive cells within the spiral ganglion was comparable between control and Sox2 cKO embryos. Apoptotic-cell number was significantly increased in the mutant by E13.5 and increased further at E14.5 (Fig. 2D and E;). Importantly, the reduction in peripheral neurite extension was already evident at E12.5, before a significant increase in apoptosis could be detected. Thus, the peripheral projection defect precedes the overt survival phenotype. Together, the lineage-tracing, proliferation and apoptosis data support a sequence in which, in the absence of epithelial Sox2, neuroblasts are generated and delaminate but subsequently show impaired peripheral projection and progressive loss.

### Transcriptomic profiling identifies Sox2-dependent epithelial signals to SGNs

To identify the epithelial signals underlying the loss of peripheral projections and neuronal survival, we performed bulk RNA sequencing on E14.5 cochlea (Fig. 3A). Differential expression analysis (log₂ fold change ≥ 1.5, FDR < 0.05) identified 623 genes downregulated after Sox2 deletion. As the dissections could retain peripheral axons, glial cells or other neuronal material, this may lead to expression changes that reflect altered tissue composition rather than epithelial transcription between mutant and control tissue. To enrich for epithelial candidates, we compared the differentially expressed genes with published single-cell RNA-seq datasets to exclude genes expressed in spiral ganglion neurons or glia (Kolla et al., 2020; Sanders & Kelley, 2022; Shrestha et al., 2018) (SI Appendix S3A). This yielded a refined set of 533 epithelial-enriched differentially expressed genes.

**Fig. 3.**
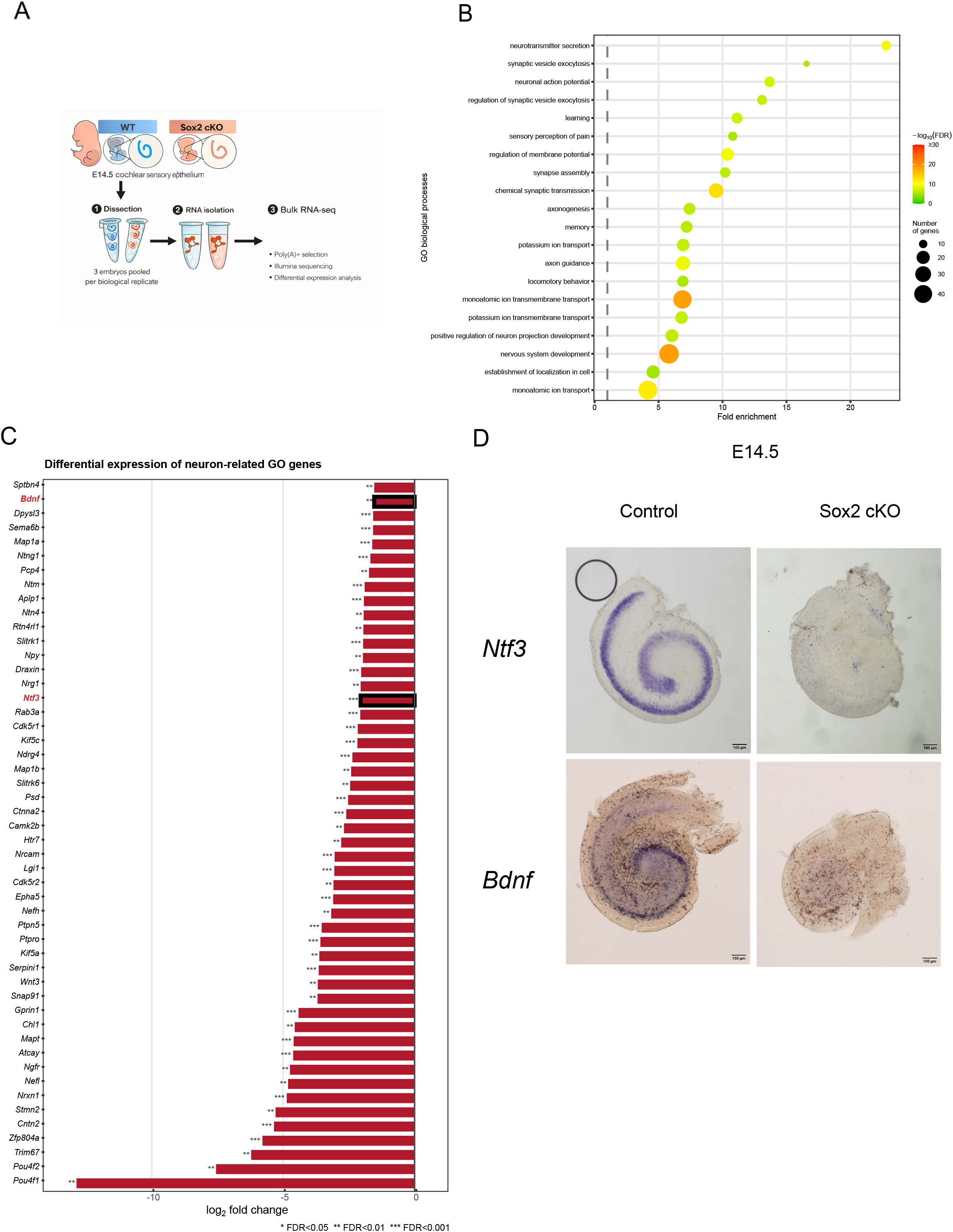
Sox2 is required for an epithelial transcriptional programme supplying neurotrophic and axon-guidance cues. (A) Experimental design. Cochlear sensory epithelium was dissected from E14.5 control and *Sox2* cKO embryos, three embryos pooled per biological replicate, and processed for poly(A)-selected bulk RNA sequencing and differential expression analysis. *n* = 3 biological replicates per genotype. (B) Gene Ontology biological process terms over-represented among the 533 epithelial-enriched downregulated genes. Dot color indicates −log₁₀(FDR), dot size the number of genes contributing to each term, and position the fold enrichment. Dashed line marks FC>1. Enrichment was assessed with DAVID -GOTERM_BP_DIRECT using the complete Mus musculus gene list as background. Significance was determined by modified Fisher exact test with Benjamini–Hochberg correction. (C) log₂ fold change (*Sox2* cKO relative to control) of the neuron-related GO genes downregulated in the *Sox2* cKO, ordered by magnitude. *Bdnf* and *Ntf3* are boxed. \**FDR* < 0.05, \*\**FDR* < 0.01, \*\*\**FDR* < 0.001. (D) Whole-mount in situ hybridization for *Ntf3* (*Upper*) and *Bdnf* (*Lower*) in E14.5 control (*Left*) and *Sox2* cKO (*Right*) cochlea. (Scale bar 100 µm.) *n* = 3 embryos per genotype per probe.

Gene Ontology analysis of the epithelially-enriched genes, down-regulated in mutant cochlea, revealed an over-representation of processes related to neuronal development and function, including chemical synaptic transmission, regulation of membrane potential, axonogenesis, axon guidance, and regulation of neuron projection development (Fig. 3B). The genes contributing to these categories were predominantly downregulated and included Slitrk1, Slitrk6, Sema6b and Ntn4, together with the neurotrophins Ntf3 and Bdnf (Fig. 3C). The reduction in Ntf3 and Bdnf was particularly notable because both factors provide target-derived support for SGN survival and peripheral axon growth and help pattern cochlear afferent innervation(Fritzsch et al., 2004; Webber & Raz, 2006).

To confirm these changes, we performed in situ hybridization for Ntf3 and Bdnf at E14.5. Both transcripts were readily detected in the control cochlear epithelium but were absent in the Sox2 cKO (Fig. 3D). Loss of Ntf3 was already evident at E12.5 (SI Appendix S3B), indicating that SOX2 is required for neurotrophin expression from the onset of peripheral axon extension. Together, these data identify a SOX2-dependent transcriptional programme that supplies neurotrophic and axon-guidance cues to the developing spiral ganglion, providing a molecular explanation for the innervation defects of the Sox2 cKO.

### Sox2 functions as activator in LGR5+ cochlear organoids

To determine how SOX2 controls the epithelial transcriptional programme identified in vivo, we required a system that provided sufficient cochlear epithelial cells for chromatin profiling. We therefore used cochlear organoids derived from LGR5^+^ supporting cells from a transgenic mouse engineered so that one copy of Lgr5 has been replaced with an inducible Cre and a GFP reporter (McLean et al., 2017). Organoids generated from Sox2^fl/fl^; Lgr5^CreERT2/+^ mice were expanded for 10 days and then treated for 5 days with either 4-hydroxytamoxifen (4-OHT) to induce Sox2 deletion or ethanol as a vehicle control (Fig. 4A). Following 4-OHT treatment, SOX2 protein was markedly reduced throughout the organoids (Fig. 4B), providing a system in which to examine the chromatin consequences of Sox2 deletion in an established cochlear epithelial population.

**Fig. 4.**
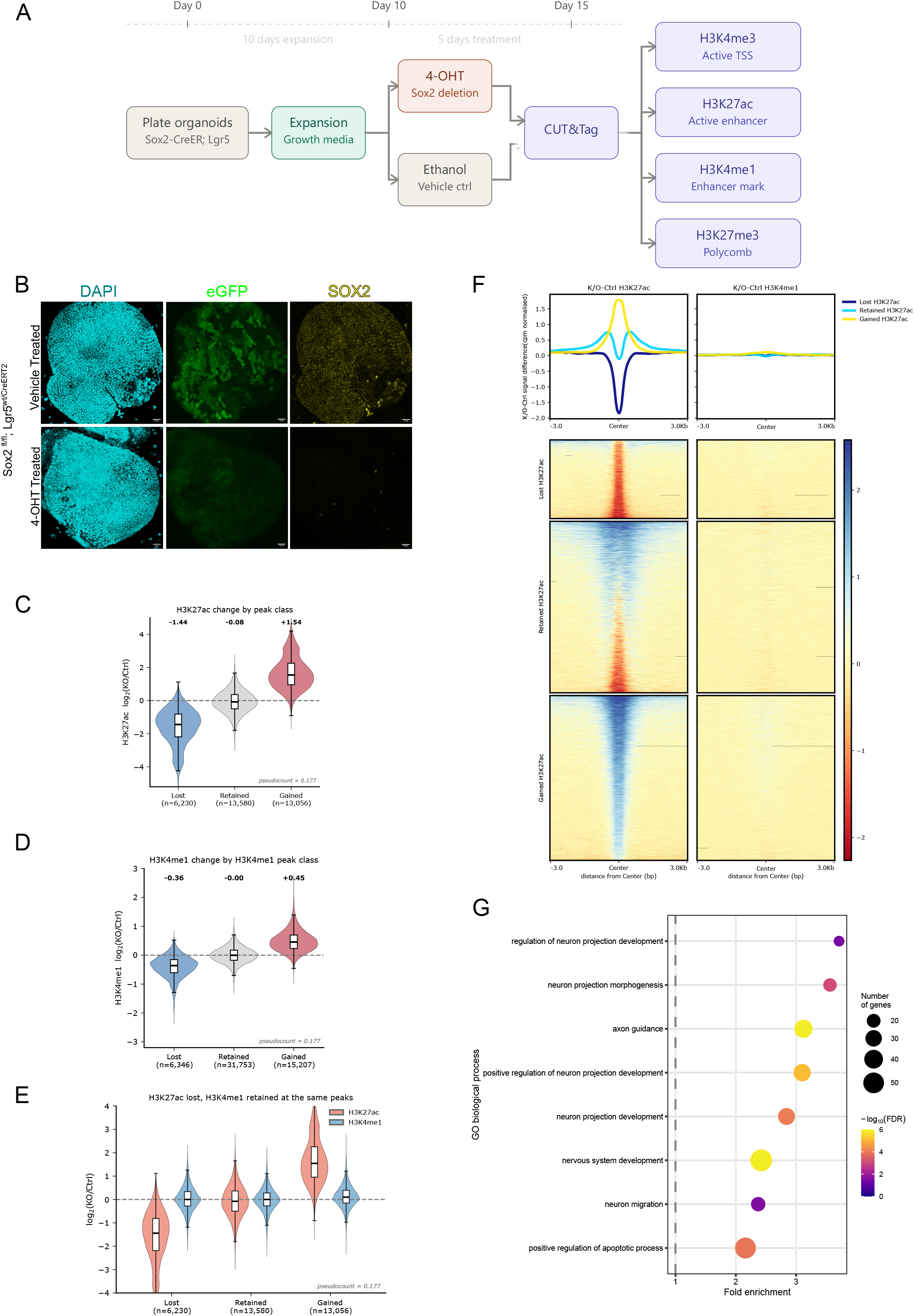
Acute Sox2 deletion in Lgr5⁺ cochlear organoids deplete H3K27ac while H3K4me1 is retained. (A) Experimental scheme. Organoids derived from *Sox2*^fl/fl^; *Lgr5*^CreERT2/+^ mice were expanded for 10 d, treated for 5 d with 4-hydroxytamoxifen (4-OHT) to delete *Sox2* or with ethanol as vehicle control, and collected at day 15 for CUT&Tag against H3K4me3, H3K27ac, H3K4me1 and H3K27me3. *n* = 1 independent organoid preparations per condition for H3K4me3, H3K27ac and H3K27me3, n=2 for H3K4me1. (B) Organoids after vehicle (*Upper*) or 4-OHT (*Lower*) treatment, labelled for DAPI, eGFP (*Lgr5* reporter) and SOX2. SOX2 is markedly reduced throughout the organoid after 4-OHT. (Scale bar 20 µm.) *n* = 3 organoids from 3 preparations per condition. (C) H3K27ac log₂(4-OHT/vehicle) at H3K27ac peaks classified as Lost, Retained or Gained. Medians are given above each class and peak numbers below. Boxes show the interquartile range, whiskers [1.5 × IQR]; ratios computed with a pseudocount of 0.177. (D) H3K4me1 log₂(4-OHT/vehicle) at H3K4me1 peaks classified independently as Lost, Retained or Gained. Note that peaks here are classified on H3K4me1 signal. (E) Paired H3K27ac and H3K4me1 log₂(4-OHT/vehicle) at the same peaks, grouped by the H3K27ac classes defined in *C*. H3K27ac falls at Lost peaks and rises at Gained peaks while H3K4me1 remains close to zero in both. (F) (*Upper*) Mean difference profiles (4-OHT minus vehicle, CPM-normalized) for H3K27ac (*Left*) and H3K4me1 (*Right*) across ±3 kb around peak centers, for the three H3K27ac classes. (*Lower*) Corresponding heatmaps, rows ordered by log2FC; color scale indicates the CPM-normalized signal difference. (paired Wilcoxon signed-rank test, rank-biserial effect size 0.97, *P* < 2.2 × 10⁻¹ ) (G) A curated Gene Ontology biological process terms over-represented among genes associated with elements that lost H3K27ac but retained H3K4me1. Dot size indicates the number of genes, color −log₁₀(FDR), and position fold enrichment; dashed line marks fold enrichment = 1. Enrichment was assessed with DAVIDGOTERM_BP_DIRECT/ALL using the complete Mus musculus gene list as background; Significance was determined by modified Fisher exact test with Benjamini–Hochberg correction.

To ask whether epithelial SOX2 establishes enhancers de novo or acts on enhancers already licensed by other factors, we performed CUT&Tag for H3K27ac and H3K4me1, together with the promoter-associated mark H3K4me3 and the Polycomb-associated mark H3K27me3. We first classified H3K27ac peaks according to their response to Sox2 deletion as Lost, Retained or Gained based on overlap. H3K27ac was strongly reduced at Lost peaks (median log fold-change, −1.44; 37% of control), remained largely unchanged at Retained peaks (−0.08; 95% of control), and increased at Gained peaks (+1.54; 292% of control; Fig. 4C). We applied the same classification to H3K4me1. Peaks classified as Lost showed reduced H3K4me1 (median log₂ fold-change −0.36; 78% of control), Retained peaks were unchanged (−0.00; 100%), and Gained peaks increased (+0.45; 137%; Fig. 4D). The magnitude of these changes was consistently smaller than for H3K27ac, consistent with H3K4me1 being the more stable of the two marks.

Enhancers that lose activation can either revert to a primed state, retaining H3K4me1 and remaining competent for reactivation, or be decommissioned (Tao et al., 2021). At peaks classified as Lost for H3K27ac, H3K4me1 showed no net change (median log₂ fold-change 0.00, Fig 4E). Comparing the two marks within individual Lost H3K27ac peak, the magnitude of H3K4me1 change was smaller than that of H3K27ac at 93% of these elements. The decoupling was therefore a property of individual enhancers rather than a population average, indicating de-activation to a primed state rather than decommissioning. Gained peaks showed the mirror image. Here H3K27ac rose sharply after *Sox2* deletion, but H3K4me1 again barely moved (+0.10; 107% of control; effect size −0.98). As H3K4me1 was already present at these elements before deletion, the data suggests that they were pre-existing enhancers that switched on. At Retained peaks, neither mark changed (effect size 0.09). Across all three classes, difference profiles confirmed that the effects of Sox2 deletion were concentrated in H3K27ac, with H3K4me1 comparatively stable on the same signal scale (Fig. 4F). SOX2 loss therefore altered enhancer activation across a largely stable primed landscape, defining a set of decoupled elements that lost H3K27ac while retaining H3K4me1.

To relate these chromatin changes to the cochlear phenotype, we asked what genes were associated with these decoupled elements. These genes were enriched for processes involved in neuron projection development, axon guidance, neuronal migration and nervous-system development (Fig. 4G). The enrichment of neuronal projection pathways is consistent with the failure of spiral ganglion peripheral axons to reach the Sox2-deficient epithelium in vivo and suggests that SOX2 maintains an epithelial regulatory programme that supports neuronal outgrowth and innervation. Together, these results show that Sox2 deletion in established cochlear organoids preferentially disrupts activation-associated H3K27ac while leaving H3K4me1 largely intact. Thus, in this cellular context, SOX2 is required to maintain the active state of regulatory elements that are already marked by H3K4me1. This pattern, in which enhancer priming persists while activation is lost, is the signature of a transcriptional activator operating at primed enhancers rather than a pioneer factor that establishes enhancers de novo.

### SOX2-dependent maintenance of an active Ntf3 enhancer

To test whether the SOX2-dependent programme identified in vivo could be reproduced in an isolated epithelial context, we measured Ntf3 expression following Sox2 deletion in cochlear organoids. Across five biological replicates, *Sox2* transcript abundance was reduced to 41% of control, whereas Ntf3 was reduced to 73% (one-sample t test on log fold-change: Sox2, P = 0.0007; Ntf3, P = 0.021; Fig. 5A). As these measurements were made in bulk cultures, residual SOX2 expression could reflect incomplete recombination or transcript persistence, potentially attenuating the measured reduction in Ntf3. Thus, NTF3 expression depends on epithelial SOX2 in both the developing cochlea and supporting-cell-derived organoids.

**Fig. 5.**
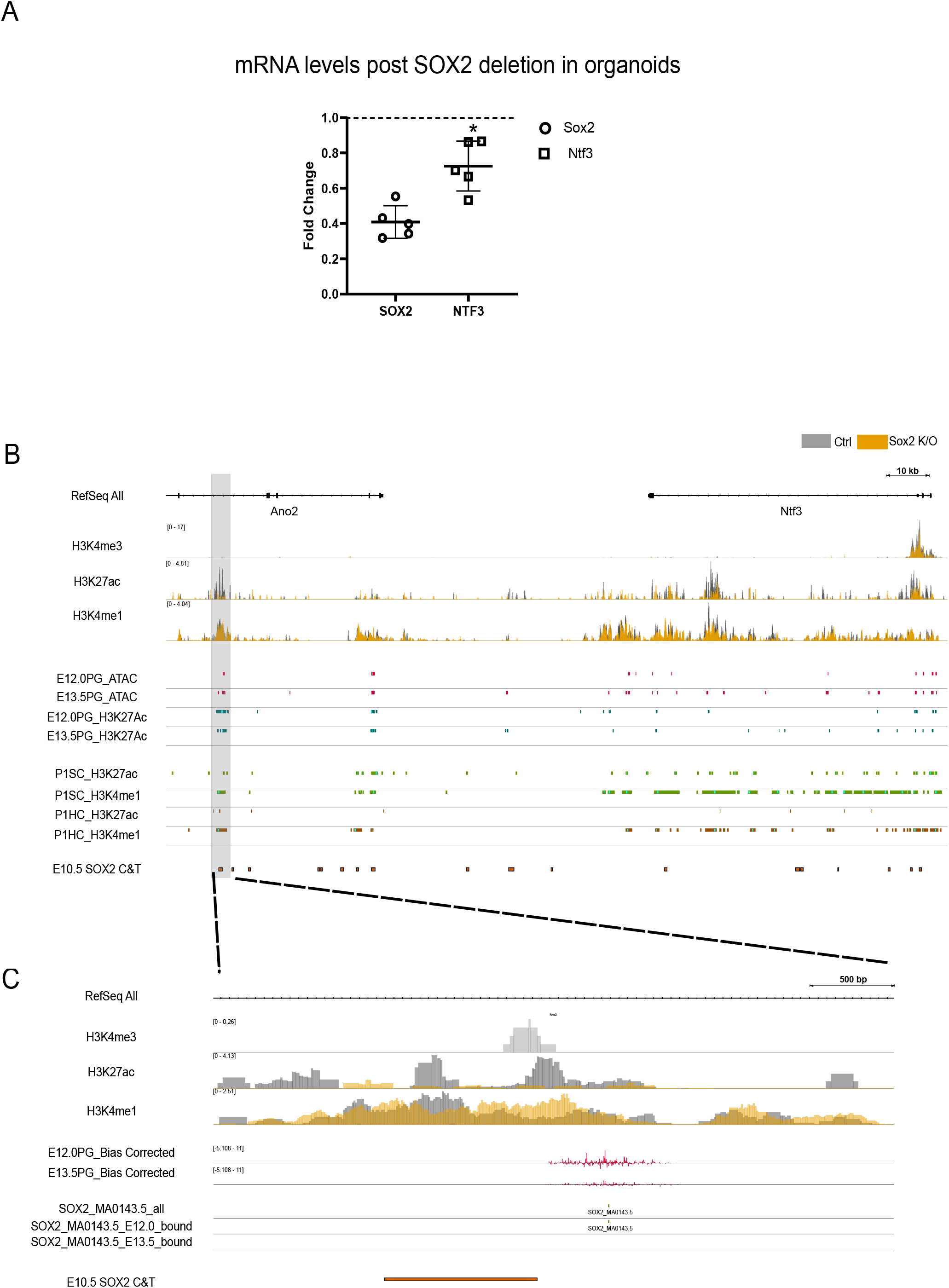
SOX2 binds and maintains H3K27ac at a distal element associated with *Ntf3*. (A) RT-qPCR for *Sox2* and *Ntf3* in cochlear organoids 5 d after 4-OHT treatment, expressed as fold change relative to vehicle-treated controls (dashed line, control = 1). Each point represents one biological replicate; bars show mean ± SD/SEM. *n* = 5. One-sample *t* test on log₂ fold change: *Sox2*, *P* = 0.0007; *Ntf3*, *P* = 0.021. (B) Genome browser view of the *Ano2*–*Ntf3* locus (mm10). (*Upper*) CUT&Tag signal for H3K4me3, H3K27ac and H3K4me1 in vehicle-treated (grey) and 4-OHT-treated (orange) organoids, overlaid on a shared [CPM-normalized] scale; data ranges are given in brackets. (*Lower*) Peak calls from published datasets: ATAC-seq and H3K27ac in sorted SOX2⁺ prosensory cells (PG) at E12.0 and E13.5; H3K27ac and H3K4me1 in P1 supporting cells (SC) and hair cells (HC); and SOX2 CUT&Tag in E10.5 otic tissue. Grey shading marks the distal element within an *Ano2* intron, ∼166 kb from the *Ntf3* promoter, expanded in *C*. (C) Expanded view of the distal element. Organoid CUT&Tag tracks as in *B*, together with TOBIAS bias-corrected ATAC signal from E12.0 and E13.5 prosensory cells, occurrences of the SOX2 motif (JASPAR MA0143.5) genome-wide and those called bound at each stage, and the overlapping E10.5 SOX2 CUT&Tag peak (orange bar). Footprint score at E12.0, 1.48 (92nd percentile of the genome-wide distribution). Footprinting performed with TOBIAS 0.17.3.

We next examined the regulatory landscape surrounding *Ntf3*. In control organoids, three candidate regulatory regions were enriched for H3K27ac: two elements within *Ntf3* introns and a distal element located approximately 166 kb from the *Ntf3* promoter within an intron of *Ano2* (Fig. 5B). Following *Sox2* deletion, H3K27ac was reduced at one intronic element and at the distal *Ano2* element, whereas the second intronic element retained H3K27ac. At both SOX2-responsive elements, H3K4me1 was comparatively stable, recapitulating at the *Ntf3* locus the genome-wide separation between loss of H3K27ac and retention of H3K4me1 (Figs. 4G, H and 5B, C). Both de-activated elements were also H3K27ac-marked in SOX2⁺ prosensory cells at E12 and E13.5 in vivo (Gnedeva et al., 2020) and the distal enhancer carried both H3K27ac and H3K4me1 in postnatal cochlear supporting cells(Tao et al., 2021), confirming that these are active enhancers used in the prosensory and supporting-cell lineages rather than organoid-specific features.

To determine whether SOX2 could act locally at these elements, we performed digital genomic foot printing with TOBIAS using ATAC-seq data(Bentsen et al., 2020) from sorted SOX2□ cochlear epithelium at E12.0 and E13.5(Gnedeva et al., 2020). At the distal *Ano2* element, a SOX2-family motif displayed a protected footprint at E12.0 (footprint score, 1.48; 92nd percentile of the genome-wide distribution), consistent with occupancy at the stage when spiral ganglion peripheral axons begin extending towards the cochlear epithelium (Fig. 5C). Coverage over this element was substantially lower at E13.5 than at E12.0 despite greater overall sequencing depth in the E13.5 libraries, precluding a reliable footprint call at the later stage. We next examined SOX2 CUT&Tag data from E10.5 otic tissue(Gao et al., 2024) as an independent, factor-specific measure of binding. A SOX2 peak overlapped the footprinted distal *Ano2* element (Fig. 5 B and C), indicating that SOX2 can engage this sequence before prosensory differentiation. Together with the loss of H3K27ac and reduction in *Ntf3* expression following Sox2 deletion, these findings support a model in which SOX2 binds a pre-marked distal regulatory element associated with *Ntf3* activation.

### *Bdnf* shows a poised, lineage-dependent mode of SOX2 regulation

During normal cochlear development, BDNF is initially expressed in cochlear progenitors and later becomes restricted to HCs (Fariñas et al., 2001). In SC-derived organoids, in which *Bdnf* is transcriptionally inactive, CUT&Tag revealed broad H3K27me3 across the *Bdnf* gene body and H3K4me3 at two positions corresponding to the start sites of the longer and shorter transcripts, but no H3K27ac peak met our calling threshold (Fig. 6A). As these marks were measured in bulk cultures, their coexistence could reflect a poised or bivalent configuration within individual cells or a mixture of states across the culture. Despite the absence of *Bdnf* transcription, and the presence of repressed marks, H3K4me1 was detected at four candidate regulatory regions: two promoter-proximal elements, associated with the long and short transcripts, and two distal elements, hereafter D1 and D2, located approximately 171kb and 156 kb from the *Bdnf* promoter. This suggests that the *Bdnf* locus therefore retains enhancer-associated chromatin in supporting cells despite being transcriptionally inactive.

**Fig. 6.**
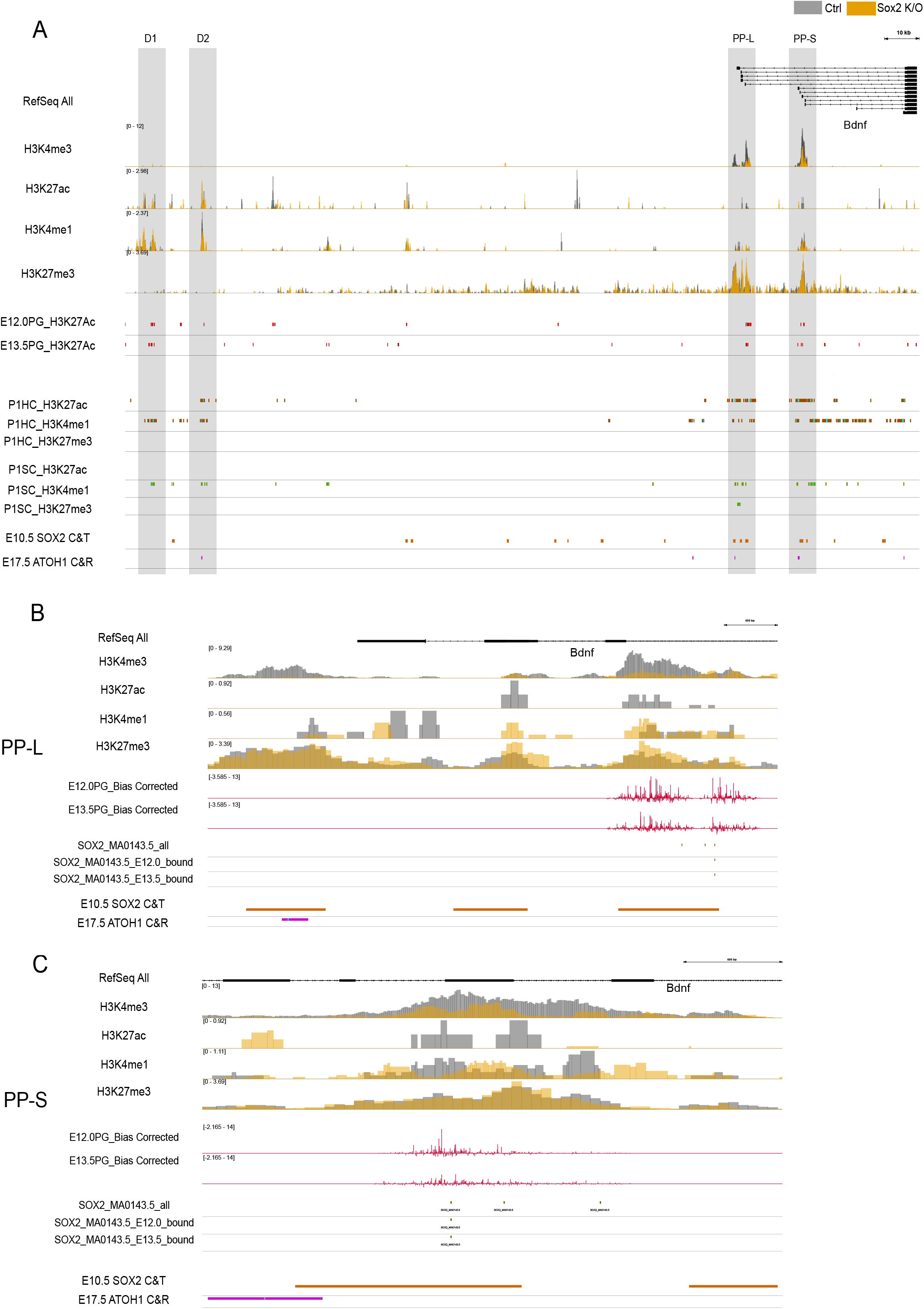
The Bdnf locus is held in a repressed but primed state in supporting-cell-derived organoids and is engaged by SOX2 and ATOH1 at different elements. (A) Genome browser view of the Bdnf locus (mm10). (Upper) CUT&Tag signal for H3K4me3, H3K27ac, H3K4me1 and H3K27me3 in vehicle-treated (grey) and 4-OHT-treated (orange) organoids, on shared [CPM-normalized] scales; data ranges in brackets. (Lower) Peak calls from published datasets: H3K27ac in sorted SOX2⁺ prosensory cells (PG) at E12.0 and E13.5; H3K27ac, H3K4me1 and H3K27me3 in P1 hair cells (HC) and supporting cells (SC); SOX2 CUT&Tag in E10.5 otic tissue; and ATOH1 CUT&RUN at E17.5. Shading marks the two distal elements D1 and D2, approximately 171 and 156 kb from the *Bdnf* promoter, and the promoter-proximal elements associated with the longer (PP-L) and shorter (PP-S) transcripts. (B and C) Expanded views of PP-L (B) and PP-S (C). Organoid CUT&Tag tracks as in A, together with TOBIAS bias-corrected ATAC signal from E12.0 and E13.5 prosensory cells, occurrences of the SOX2 motif (JASPAR MA0143.5) genome-wide and those called bound at each stage, the E10.5 SOX2 CUT&Tag peak (orange) and the E17.5 ATOH1 CUT&RUN peak (magenta).

We thus asked whether these H3K4me1-marked elements had been active earlier, when *Bdnf* was expressed broadly in cochlear progenitors. Comparison across embryonic and postnatal datasets revealed that the four elements followed distinct developmental trajectories (Fig. 6). In embryonic SOX2□ cochlear epithelium, both promoter-proximal elements carried H3K27ac. D1 was also acetylated at E12.0 and E13.5, whereas D2 was acetylated at E12.0 only, consistent with the broader expression of *Bdnf* in cochlear progenitors. Postnatally, D1 retained H3K4me1 but lacked H3K27ac in both hair cells and supporting cells. By contrast, D2 and the two promoter-proximal elements carried H3K27ac in hair cells, where BDNF is expressed, but retained H3K4me1 without H3K27ac in supporting cells(Tao et al., 2021). This suggests that regulatory competence persists in both epithelial descendants, whereas activation becomes restricted to selected elements in the hair-cell lineage. However, the embryonic SOX2 population also includes Kölliker’s organ and adjacent non-sensory epithelium, so its H3K27ac signal cannot be assigned exclusively to prosensory cells.

We next asked whether SOX2 occupies these elements. Of seven SOX2 motif matches within the accessible *Bdnf* 5′ cluster, SOX2 was footprinted at both promoter proximal elements at both stages: chr2:109,678,048–109,678,064 (E12.0, 1.78, 94th percentile; E13.5, 1.84, 94th percentile) and chr2:109,678,130–109,678,146 (E12.0, 2.34, 97th percentile; E13.5, 1.40, 90th percentile); the remaining matches were not classified as bound under the same criteria. SOX2 CUT&Tag from E10.5 otic tissue independently identified peaks overlapping both promoter-proximal elements, providing factor-specific evidence that SOX2 engages the Bdnf 5′ region before lineage differentiation. No SOX2 footprint was detected at either distal element at E12.0 or E13.5(SI Appendix S4A and S4B). A single SOX2 CUT&Tag call lay close to D1, but it was not supported by a footprint at either stage and, as the SOX2 CUT&Tag dataset comprises a single replicate, we did not consider it further. H3K4me1 at the promoter-proximal elements was retained following Sox2 deletion in organoids, indicating that ongoing SOX2 activity is not required for short-term maintenance of this mark in an established epithelial population.

The pattern of occupancy changed after differentiation. ATOH1 CUT&RUN from E17.5 hair cells identified binding at both promoter-proximal elements and at D2, which acquires H3K27ac in postnatal hair cells; ATOH1 did not bind D1(Yu et al., 2021)(SI Appendix S4A). Thus, SOX2 and ATOH1 engage the Bdnf locus at different stages and with different distributions. SOX2 binding is confined to the promoter-proximal elements and is detected in otic tissue and in SOX2⁺ epithelial progenitors, whereas ATOH1 occupies those same elements in addition to a distal element that SOX2 does not engage, after hair-cell identity has been acquired.

Together, *Ntf3* and *Bdnf* illustrate complementary modes of neurotrophin regulation. At *Ntf3*, SOX2 occupies the responsive enhancer directly and is required to maintain its H3K27ac, while H3K4me1 persists in its absence. At *Bdnf*, SOX2 occupies promoter-proximal, H3K4me1-marked elements within a transcriptionally repressed locus, whereas the element that acquires hair-cell-specific H3K27ac and ATOH1 binding lies distal and is not engaged by SOX2 at any stage examined. These findings support a locus-level regulatory transition in which *Bdnf* is interpreted by distinct transcription-factor inputs as cochlear epithelial lineages diverge.

## DISCUSSION

Our findings identify SOX2 as a regulator of signalling between the cochlear sensory epithelium and spiral ganglion neurons. Following Sox2 deletion, we find a reduction of SGN peripheral projections before we observe increased neuronal apoptosis. However, neuroblast generation, delamination and proliferation were largely preserved, suggesting that neuroblast specification occurs normally. This is consistent with the timing of Emx2-Cre activation, which drives the deletion of Sox2 after its requirement for neurogenesis. Moreover, our data suggest that peripheral axons must reconnect with the differentiating epithelium to engage with epithelial signals for survival. Consistent with this, transcriptomic profiling identified a SOX2-dependent epithelial programme enriched for neurotrophic factors, axon-guidance molecules and regulators of neuronal projection development.

The loss of *Ntf3* and *Bdnf* links the epithelial defect and the SGN phenotype. These neurotrophins occupy complementary spatial and cellular domains during cochlear development and together support SGN survival and afferent innervation(Fariñas et al., 2001; Fritzsch et al., 2004; Pirvola et al., 1992). *Ntf3* becomes concentrated largely in supporting and other non-hair cells, whereas *Bdnf* is associated with differentiating hair cells(Fariñas et al., 2001). Their loss in the Sox2 cKO is therefore consistent with the failure of peripheral innervation followed by progressive neuronal death. However, Sox2 deletion also reduced other neuronal-support and guidance genes, including *Slitrk1*, *Slitrk6*, *Sema6b* and *Ntn4*. Amongst these, *Slitrk6* has been well characterised in the developing cochlea, showing expression early in the otocyst and later, strongly in supporting cells and weakly in hair cells. Mouse mutants of Slitrk6 show reduced cochlea innervation(Katayama et al., 2009). Thus, *Ntf3* and *Bdnf* can be considered exemplars of a broader SOX2-dependent programme rather than the sole causes of the phenotype.

The chromatin analysis suggests how SOX2 sustains this programme. Sox2 deletion in cochlear organoids produced pronounced changes in H3K27ac at a subset of regulatory elements, whereas H3K4me1 remained stable in the same regions. We concluded that SOX2 is required to maintain activation-associated acetylation at elements that are already marked by H3K4me1. Elements showing this separation between H3K27ac and H3K4me1 were associated with genes involved in axon guidance and neuronal projection development, connecting the chromatin response to the innervation phenotype. We note that Sox2 was deleted after organoid expansion, so these experiments do not establish that enhancer marking was originally acquired independently of SOX2 or exclude an earlier pioneer function. Rather, they support a requirement for SOX2 in maintaining enhancer activity in an established cochlear epithelial state. They do not, however, show that SOX2 is dispensable for the initial deposition of H3K4me1, because Sox2 was deleted only after organoid expansion. H3K4me1 is deposited by MLL3 and MLL4, which are recruited by DNA-binding factors rather than binding DNA directly(Hu et al., 2013). At elements occupied by several transcription factors, loss of one may abolish activation while the primed state is sustained by the others. This possibility is important to consider as in the cochlear epithelium, SOX9 and SOX21 are also expressed in supporting cells (Hosoya et al., 2011; Mak et al., 2009)and recognise HMG motifs of the same class. Thus, footprints at SOX-family motifs may therefore indicate occupancy by the family rather than by SOX2 specifically. Factor-specific occupancy data and combinatorial perturbation of these SOX proteins, will be required to determine how enhancer marking and activation are established and maintained.

*Ntf3* illustrates the maintenance of a lineage-appropriate active programme. *Sox2* deletion reduced *Ntf3* transcription and H3K27ac at intronic and distal candidate regulatory elements while H3K4me1 persisted. The distal element within Ano2 was also marked in embryonic SOX2□ cochlear epithelium and postnatal supporting cells. An E12.0 SOX-family footprint and an overlapping E10.5 SOX2 CUT&Tag peak provide complementary evidence that SOX2 can engage this element during otic development. These findings identify it as a strong candidate through which SOX2 contributes to *Ntf*3 activation, although targeted perturbation will be required to establish its regulatory relationship with *Ntf3*.

*Bdnf* displayed a contrasting configuration. In supporting-cell-derived organoids, where *Bdnf* was transcriptionally inactive, the locus retained H3K4me1 at four candidate regulatory regions but lacked robust H3K27ac. Two of these lie promoter-proximal, associated with the longer and shorter transcripts, and two lie distal. All four carried H3K27ac in embryonic SOX2⁺ epithelium, at stages when *Bdnf* is transcribed in progenitors. Postnatally they diverged. One distal element retained H3K4me1 alone in both hair cells and supporting cells, whereas the other acquired H3K27ac in hair cells while remaining H3K4me1-marked in supporting cells. SOX2 occupied both promoter-proximal elements in progenitors. ATOH1 later occupied these same elements together with the distal element that acquires hair-cell H3K27ac, which SOX2 did not engage at any stage examined. The data therefore do not indicate replacement of SOX2 by ATOH1 at a single enhancer. Instead, they support a locus-level regulatory transition in which activation is distributed across the locus in progenitors and subsequently retained at a subset of elements in the hair-cell lineage. *Ntf3* and *Bdnf* thus represent complementary outcomes of lineage divergence: maintenance of an active supporting-cell-associated programme and redistribution of regulatory inputs during hair-cell-associated gene activation.

This pattern is not unique to the cochlea. Lineage-dependent reinterpretation of inherited enhancers has been described during neural, endodermal, haematopoietic and muscle differentiation, where regulatory elements established in a progenitor state are selectively retained, activated or decommissioned as cells become restricted(Aksoy et al., 2013; Bergsland et al., 2011; Lilja et al., 2017). In those settings, the mechanism partitions cell-intrinsic identity programmes among descendants. The cochlea suggests a further use: the inherited landscape sustains expression of secreted factors that serve a different lineage altogether, so that regulatory memory in the epithelium underwrites the survival and wiring of neurons that separated from it days earlier.

Several limitations qualify this model. Although the timing of the projection defect and preservation of neuroblast proliferation support an epithelial contribution, lineage-reporter labelling within the ganglion means that a direct effect of *Sox2* deletion in a subset of neuronal or glial cells cannot be excluded. Moreover, supporting-cell-derived organoids do not fully reproduce the embryonic prosensory state, and the locus comparisons integrate bulk datasets from different stages and cell populations. Finally, chromatin marking and genomic proximity do not demonstrate enhancer function or enhancer–gene assignment. Epithelial-restricted deletion, neurotrophin rescue and targeted perturbation of the candidate regulatory elements will be required to test these relationships.

Together, our findings extend the role of SOX2 from sensory fate specification to the coordination of lineages with a shared otic ontogeny. Neuroblasts and epithelial progenitors arise from a common otic territory and follow divergent trajectories, yet must remain functionally connected as SGN axons return to, and depend upon, the sensory epithelium. Our data suggest that SOX2 preserves this relationship by maintaining activation-associated acetylation at H3K4me1-marked epithelial enhancers, thereby coupling epithelial identity to neuronal-support signals. *Ntf3* and *Bdnf* illustrate how this regulatory competence can be interpreted differently along supporting-cell and hair-cell trajectories: an inherited landscape whose output is partitioned rather than erased as descendants acquire distinct identities.

More broadly, our findings extend the role of SOX2 beyond sensory fate specification. Neuroblasts and sensory epithelial cells arise from a common otic territory but become physically separated before SGN axons reconnect with their epithelial targets. By maintaining an epithelial regulatory programme that includes neuronal-support signals, SOX2 helps coordinate the development of these diverging populations. The cochlea therefore provides an example in which lineage-dependent interpretation of an inherited regulatory landscape may preserve communication between descendants of a common progenitor even as they acquire distinct identities.

## MATERIALS AND METHODS

### Mice

The following mouse strains were used in this study. B6.Cg-Emx2^tm2^ ^(cre)Sia/SiaRbrc^ mice (RIKEN BRC, RBRC04895; RRID: *RBRC04895*) carry a Cre recombinase knock-in allele at the *Emx2* locus, enabling Emx2-driven Cre activity in the developing inner ear and associated neuroepithelial tissues. B6.129P2-Lgr5^tm1^ ^(cre/ERT2)Cle/J^ mice (The Jackson Laboratory, Stock No. 008875; RRID: *IMSR_JAX:008875*) express a tamoxifen-inducible CreERT2 fusion protein from the *Lgr5* locus, permitting temporal control of recombination in Lgr5-positive supporting cells. **Sox2^tm1.1Lan/J^** mice (The Jackson Laboratory, Stock No. 013093; RRID: *IMSR_JAX:013093*) harbour a floxed *Sox2* allele that enables conditional *Sox2* deletion upon Cre activation. All strains were maintained on a C57BL/6J background and intercrossed as required to generate the experimental genotypes.

### Tissue Dissection and Fixation

Inner ears from staged mouse embryos were dissected in ice-cold phosphate-buffered saline (PBS) using fine forceps. Tissues were fixed in 4% paraformaldehyde (PFA) for 4 h at room temperature (RT), followed by three washes in PBS (15 min each).

### Whole-mount Immunostaining of Sensory Epithelia

Following fixation, inner ears were further dissected under a stereomicroscope to expose the sensory epithelia. Tissues were permeabilized in 0.5% Triton X-100 in PBS for 30 min at RT and subsequently blocked for 1 h at RT in blocking solution (10% heat-inactivated goat serum, 1% bovine serum albumin, 0.5% Triton X-100 in PBS).

Sensory epithelia were incubated with primary antibodies (Table 1) diluted in blocking solution overnight at 4°C. The next day, samples were washed for 4 h in wash buffer (0.5% Triton X-100 in PBS), with buffer exchanged every 30 min. Tissues were then incubated with Alexa Fluor–conjugated secondary antibodies and Alexa Fluor– conjugated phalloidin for 1 h at RT. After secondary incubation, samples were washed for 3 h in wash buffer, with solution changed every 30 min. Sensory epithelia were mounted on glass slides using an aqueous mounting medium.

**Table 1:** Antibodies Used

| Antibody | Company | Cat No- |
| --- | --- | --- |
| SOX2 | Invitrogen | 14-9811-82 |
| BETA-SPECTRIN2 | BD Biosciences | 612563 |
| PROX1 | Merck Millipore | AB5475 |
| beta3-Tubulin (TUJ1) | Synaptic Systems | 302 306 |
| NeuN | Abcam | ab177487 |
| Cleaved CASPASE 3 | Cell Signalling Technology | 9664S |
| NEUROD1 | Cell Signalling Technology | 62953S |

### Cryosectioning and Immunostaining

For sectioning, fixed inner ears were washed in PBS and cryoprotected by incubation in a sucrose gradient (10%, 20%, and 30% sucrose in PBS). Tissues were embedded in OCT compound, frozen, and sectioned at 20 μm thickness using a cryostat. Sections were collected on charged slides and processed for immunostaining following the procedure described for whole-mount samples, beginning with permeabilization.

### Confocal Microscopy

Whole Mount epithelia and sections were imaged on an Olympus FluoView 3000 inverted confocal microscope controlled by FV31S-SW software (Olympus, Japan) at the Central Imaging and Flow Cytometry Facility (CIFF), NCBS. Z-stacks were collected with a step size of 0.5 µm, and xy pixel size was set according to Nyquist sampling criteria.

### Paint Fill

The paint-filling procedure was performed following a protocol provided by Dr. Doris Wu(Morsli et al., 1998). Briefly, heads from E14.5 mouse embryos were fixed overnight in Bodian’s fixative and subsequently dehydrated in 100% ethanol for four days to ensure complete dehydration. Samples were cleared in methyl salicylate until transparent. Cleared heads were bisected along the midline, and the brain was removed to expose the inner ear. White enamel paint, diluted in methyl salicylate, was introduced into the mid-turn of the organ of Corti using a fine glass capillary.

### EdU Click Reaction

Timed-pregnant females were administered 5-ethynyl-2’-deoxyuridine (EdU; Sigma, 900584) diluted in sterile PBS at a dose of 2 mg per 40 g body weight for 6hrs on the indicated time points. Embryonic tissues were collected at the indicated time points and processed for EdU detection. The Click-iT EdU reaction was performed according to the manufacturer’s instructions (Invitrogen C10337), ensuring complete protection from light. Following the click reaction, samples were subjected to standard immunostaining procedures using the appropriate primary and secondary antibodies.

### In Situ Hybridisation

All solutions were rendered RNase-free by treatment with diethyl pyrocarbonate (DEPC). Solutions were incubated under standard conditions to inactivate RNases, and all reagents not compatible with autoclaving were prepared using DEPC-treated water to maintain RNase-free conditions throughout the procedure.

Mouse embryo heads (E15.5) were fixed in 4% paraformaldehyde (PFA) and subsequently washed thoroughly in DEPC-treated PBS (DEPC–PBS). Cochleae were dissected and dehydrated through a graded methanol/PBS series before being washed in 100% methanol and stored at –20°C.

Cochleae were gradually rehydrated into PBST (PBS with detergent) on ice and post-fixed using a PFA–glutaraldehyde mixture. After PBST washes, samples were equilibrated in pre-hybridisation solution and incubated under standard pre-hybridisation conditions. Digoxigenin (DIG)-labelled RNA probes were added at a typical working concentration, and hybridisation was performed overnight at elevated temperature.

Following hybridisation, samples underwent high-stringency washes at maintained temperature, followed by washes in maleic acid buffer (MABT). Non-specific binding was blocked using a solution containing commercial blocking reagent and heat-inactivated goat serum. Cochleae were then incubated with anti-DIG alkaline phosphatase–conjugated antibody under standard conditions. Extensive washing in MABT was performed to reduce background signal. Colour development was carried out using either NBT/BCIP in NTMT buffer or BM-Purple (Roche) until sufficient staining was obtained. Cochleae were equilibrated in 60% glycerol for optical clearing and imaged using a stereomicroscope. DIG-labelled RNA probes were synthesised using Roche transcription reagents following the manufacturer’s instructions.

### Organoid Culture

Cochlear organoids were generated following established protocols for clonal expansion of Lgr5⁺ epithelial progenitors(McLean et al., 2017). Briefly, cochlear epithelia were micro-dissected in HBSS, and the sensory epithelium (organ of Corti) was isolated from the surrounding mesenchyme and stria vascularis. Tissue was incubated in Cell Recovery Solution for 1 h at 4 °C to separate the epithelial sheet, followed by enzymatic dissociation in TrypLE Express for 15–20 min at 37 °C with gentle trituration to obtain a single-cell suspension. Cells were filtered (40 µm) and seeded as single cells into 3D Matrigel droplets, optionally supplemented with laminin (0.015 mg/mL) to enhance epithelial attachment. Organoids were cultured in serum-free DMEM/F12 (1:1) containing GlutaMAX, N2 and B27 supplements, and a growth factor/small-molecule cocktail (EGF 50 ng/mL, bFGF 50 ng/mL, IGF-1 50 ng/mL, CHIR99021 3 µM, valproic acid 1 mM, 2-phospho-L-ascorbic acid 100 µg/mL, and the ALK5 inhibitor 616452 at 2 µM), as previously described. Media were changed every other day. For Sox2 deletion experiments, 4-hydroxytamoxifen (25 ng/mL) was added at day 10. After 20 days of expansion, organoids were harvested for downstream applications including immunostaining, RNA extraction, and CUT&Tag profiling.

### RT-qPCR

Complementary DNA was synthesised from 1 ug total RNA using Takara PrimeScript™ 1st strand cDNA Synthesis Kit (CatNo6110A) with random hexamers, according to the manufacturer’s instructions. Quantitative PCR was performed on Applied Biosystems ViiA 7 Real-Time PCR System using Applied Biosystems SYBR Green Universal Master Mix, with 3 technical replicates per biological sample. Relative expression was calculated by the 2^(−ΔΔCt)^ method, normalised to Gapdh and expressed relative to vehicle-treated controls. Primer sequences are listed in Table 2.

**Table 2:**
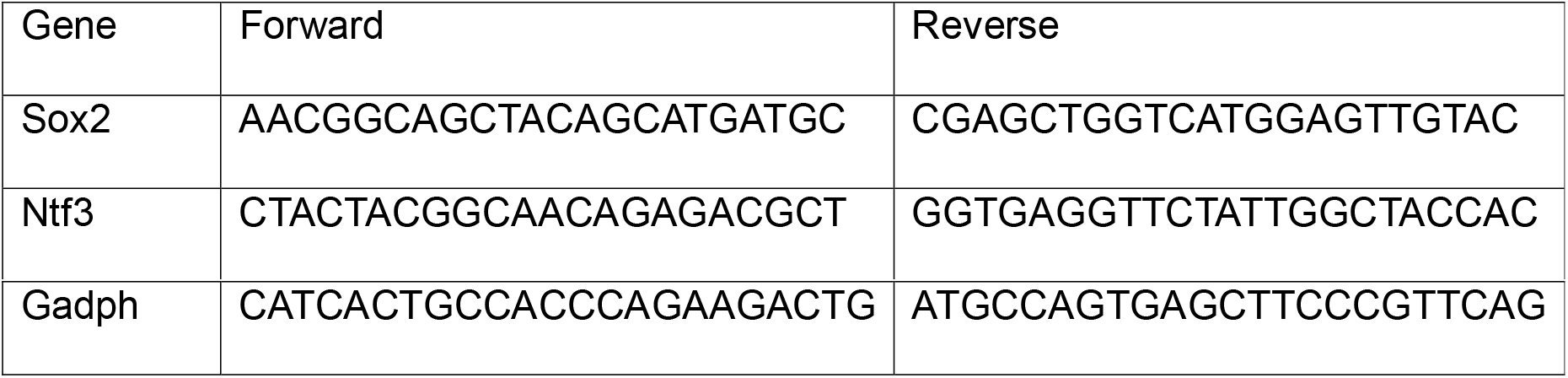
qPCR Primers

### RNA-seq + analysis pipeline

Organ of Corti from E14.5 embryo were dissected in HBSS and collected in 200ul of Trizol. Tail clipping from the embryo was used to genotype. Organ of Corti from three genotypically same embryos were pooled and treated as biological sample. Total RNA was manually isolated followed by polyA based enrichment using NEBNext Poly (A) mRNA Magnetic Isolation Module. NEBNext® Ultra™ II Directional RNA Library Preparation kit was used for library preparation.

Raw FASTQ reads (150 bp, paired-end) for both wild-type and SOX2 mutant mouse samples were generated using the Illumina Novaseq 6000 platform. Read quality was assessed using FastQC (version 0.12.1). High-quality reads were then aligned to the mouse reference genome (mm10) using HISAT2 version 2.2.1 (Kim et al., 2019), producing SAM files. These files were subsequently sorted and converted to BAM format using Samtools (version 1.22.1).

Transcript assembly and quantification were performed with StringTie version 2.2.1(Pertea et al., 2015). Transcript assembly enables reconstruction of full-length transcript isoforms, detection of alternative splicing events, identification of novel transcripts, and improved transcript-level quantification. Assemblies were generated for each sample independently and then combined using the StringTie–merge function to obtain a unified set of transcript models consistent across samples. Final transcript abundances were then re-estimated by running StringTie against the merged annotation. An in-house Python script was used to process StringTie output and generate the final gene count matrix.

Differential gene expression analysis and replicate clustering (MDS plot) were performed using the edgeR (version 4.0.2) (M. D. Robinson et al., 2010)package in R (version 4.3.3). Genes with more than 10 reads in at least two samples were retained for analysis. Library size and composition normalization was performed using the calcNormFactors () function (TMM method). Differential expression was assessed using the makeContrasts () function. Genes with a log2 fold change ≥ 1.5 and an adjusted p-value < 0.05 were considered significantly differentially expressed. These differently expressed genes (DEGs) were then compared to scRNA datasets for SGN and glia(Sanders & Kelley, 2022), and Organ of Corti(Kolla et al., 2020) to exclude genes only present in the SGN and Glial and a list of refined DEGs.

### Gene Ontology Analysis

Genes associated with differential H3K27ac peaks were assigned by proximity using ChIPseeker, and the resulting gene lists were analysed for functional enrichment using DAVID (v6.8/2021 update;(Huang et al., 2009)). Enrichment was assessed against the Mus musculus genome background, and GOTERM_BP_DIRECT was used to report biological process terms. Terms were considered significant at a Benjamini–Hochberg-corrected FDR < 0.05.

### CUT and Tag and Analysis

CUT&Tag(Kaya-Okur et al., 2019) was performed using the CUT&Tag Assay Kit (Cell Signaling Technology, CST) following the manufacturer’s protocol with minor modifications. Briefly, 100,000 cells from dissociated organoids were immobilized on Concanavalin A–coated magnetic beads and permeabilized in the supplied Dig-Wash buffer. Samples were incubated overnight at 4 °C with primary antibodies against the target of interest, followed by a 1-hour incubation with CST’s secondary anti-rabbit IgG–Tn5 transposome complex. After extensive washes to remove unbound enzyme, tagmentation was activated by adding Mg²⁺ at 37 °C for 1 hour to generate indexed DNA fragments at protein–DNA binding sites.Reactions were stopped using the CST stop buffer, and DNA fragments were released by heating at 58 °C for overnight.

Libraries were amplified using the CST indexing primers for 12–15 PCR cycles, and amplified fragments were purified using SPRIselect beads (Beckman Coulter). Library size distribution was assessed by Bioanalyzer, and libraries were sequenced on an Illumina platform using paired-end 150bp reads. Sequencing reads were trimmed, aligned to the mm10 genome using Bowtie2(Langmead & Salzberg, 2012) with default parameters, and peaks were called using MACS2 (Zhang et al., 2008)(narrow peaks for H3K4me3 and broad peaks for H3K27ac,H3K4me1 and H3K27me3). Bigwig files for visualisation were generated using bamCompare(Ramírez et al., 2016) with cpm as the normalisation. IGV 2.19.7(J. T. Robinson et al., 2011) was used for visualisation.

CUT&Tag was performed on a single biological replicate per condition for each histone modification apart from H3K4me1 which has two replicates. Peaks were therefore classified by overlap rather than by statistical differential binding: peaks called in vehicle-treated but not 4-OHT-treated organoids were designated Lost, those called in 4-OHT-treated but not vehicle-treated organoids Gained, and those called in both Retained (bedtools intersect). Per-peak signal changes were quantified as log₂((KO + c)/(Ctrl + c)) from mean CPM over each peak interval, with a fixed pseudocount c = 0.177.

### Genomic footprinting

ATAC-seq from sorted SOX2⁺ cochlear progenitors at E12.0 and E13.5 (Gnedeva et al., 2020)was analysed against mm10. Reads were aligned with Bowtie2 and filtered to properly paired, uniquely mapping, non-duplicate reads, excluding mitochondrial reads and the ENCODE mm10 blacklist (v2). Alignments were not Tn5-shifted, as TOBIAS applies the +4/−5 offset internally. Replicates were merged per stage, and a consensus peak set defined as regions called in at least two of three replicates (bedtools merge).

Footprinting was performed with TOBIAS 0.17.3(Bentsen et al., 2020),Tn5 bias correction with ATACorrect, per-base footprint scoring with ScoreBigwig, and occupancy calling with BINDetect, run once across both conditions so that thresholds were applied identically to E12.0 and E13.5. Motifs for the genome-wide survey of co-occupying factors were occupied from JASPAR CORE(Castro-Mondragon et al., 2022) vertebrates non-redundant. Default thresholds were used (--motif_pvalue [1e-4], -- bound_pvalue [0.001]) with no locus- or condition-specific adjustment; footprint scores at individual sites are reported as percentiles of the genome-wide distribution for that motif. Because footprint detection is limited by local read depth, coverage at individual elements was quantified with samtools and compared between stages; elements with substantially unequal coverage were reported as indeterminate rather than differentially occupied.

### Statistics

Statistical analyses were performed in R (v 4.6.0) and GraphPad Prism (v 8.0.2). For imaging and expression analyses, *n* refers to the number of biological replicates (individual embryos or independent organoid preparations), as stated in each figure legend. Comparisons between two groups were made using Student’s *t*-test, or a one-sample *t*-test where values were expressed relative to a normalised control. Data are presented as mean ± SD unless otherwise indicated. For genome-wide analyses, per-peak comparisons between histone marks were made using the two-sided Wilcoxon signed-rank test with paired samples, and effect sizes are reported as rank-biserial correlations; *P* values below the limits of double-precision arithmetic are reported as *P* < 2.2 × 10⁻¹□. Gene ontology enrichment was assessed using a modified Fisher’s exact test with Benjamini–Hochberg correction, and terms were considered significant at FDR < 0.05. Differential gene expression was assessed in edgeR using quasi-likelihood F-tests, with genes considered differentially expressed at FDR < 0.05 and |log₂ fold-change| ≥ 1.5. The specific test, *n* and significance threshold for each experiment are given in the corresponding figure legend.

## ACKNOWLEDGEMENTS

This work was supported by the Department of Atomic Energy, Government of India, Project Identification No. RTI 4006, and grants from the Royal National Institute for Deaf People (IPG programme), SERB and TIFR Infosys-Leading Edge Grant. We acknowledge the support of the Animal Care and Resources Centre, NGS and Central Imaging Facility at NCBS. A.P. acknowledges the Simons Foundation International for support through the Simons-Ashoka early-career fellowship. We thank Dr Gaiti Hasan, Dr Dimple Notani and Dr Jasmine Dhall for generously sharing reagents and Dr Aswin Sai Narain Seshasayee for sharing computational infrastructure. We are grateful to members of the Ear Lab for discussions.

## Author Contributions

SR and RL conceived the study and designed the experiments. SR performed the mouse genetics, immunostaining and imaging, generated and cultured the cochlear organoids, and performed the qPCR, CUT&Tag and RNA-seq experiments. SR performed the computational analyses for CUT&Tag and AD performed the computational analyses for RNA-seq. AP contributed to the immunostaining experiments. RK and SR performed the paintfill experiments. LK and SR performed the ISH experiments. PC performed the length quantification experiments. SR and RL wrote the manuscript with input from all authors. RL supervised the study and acquired funding. All authors read and approved the final manuscript.

## Competing Interests

The authors declare no competing interests.

## Data Availability

RNA-sequencing and CUT&Tag datasets generated in this study will be deposited in the NCBI Gene Expression Omnibus and accession numbers provided upon acceptance. Publicly available datasets analysed in this study are available under accessions GSE150391(Yu et al., 2021), GSE149254(Gnedeva et al., 2020),GSE239363(Gao et al., 2024) and GSE150000(Tao et al., 2021). All other data supporting the findings are available from the corresponding author on reasonable request.

## Notes

### Competing Interest Statement

The authors have declared no competing interest.

### Summary of Updates

Addition of new data to provide more confidence in analysis

