## Supplementary material for "SOX2-dependent enhancer activation couples epithelial identity to neuronal maintenance": SI

**Fig. S1. Loss of SOX2 from the cochlear epithelium disrupts prosensory identity and sensory differentiation without gross malformation of the membranous labyrinth.**

(A and B) Transverse sections through the apical, middle and basal cochlear duct at E14.5 from control (A) and Sox2 cKO (B) embryos, labelled for DAPI, F-actin, p27KIP1 and SOX2. Maximum-intensity projections of 20-μm sections. Arrows indicate the prosensory domain, which lacks SOX2 and shows reduced p27KIP1 at all three positions in the cKO. (Scale bar 20 μm.) n = 3 embryos per genotype.

(C) Whole-mount basal cochlear turn at E15.5, control (Upper) and Sox2 cKO (Lower), labelled for β-spectrin, PROX1, SOX2 and DAPI. Organized β-spectrin-positive hair cells and PROX1-positive supporting cells are absent from the cKO. (Scale bars 100 μm.) n = 3 per genotype.

(D and E) Paint-filled inner ears at E14.5, control (C) and Sox2 cKO (D). The major structures of the membranous labyrinth are present in both. (Scale bars, 200 μm.) n = 3 per genotype.

(E) Whole-mount E16.5 cochleae, control (Upper) and Sox2 cKO (Lower), labelled for DAPI, SOX2 and F-actin. (Scale bar 100 μm.)

(F) Cochlear duct length at E14.5, E16.5 and E18.5. Each point represents one cochlea; bars show mean ± SD/SEM. n = 2-5 per genotype per stage. Paired t test; ***P =0.004; the difference at E14.5 was not significant (P =0.08).

**Fig. S2. Neuroblast proliferation is unchanged in *Sox2* cKO embryos at E13.5.**

(*A*) Serial transverse sections through the E13.5 inner ear of control (*Left*) and *Sox2* cKO (*Right*) embryos, labelled for DAPI, NeuroD1, EdU and SOX2. Every fourth 20-µm section is shown, arranged posterior to anterior. (Scale bar 20µm.) *n* = 4 cochlea per genotype.

**Fig. S3. Exclusion of neuronal and glial transcripts from the *Sox2* cKO downregulated gene set, and early loss of *Ntf3*.**

(*A*) Four-way Venn diagram comparing the 623 genes significantly downregulated in the *Sox2* cKO (Sox2SigDwn) with genes detected in published single-cell RNA-seq datasets of spiral ganglion neurons (SGN scRNA), glia (Glial) and organ of Corti (OC scRNA). Genes shared with the SGN and glial datasets were excluded, yielding 533 epithelial-enriched genes.

(*B*) Whole-mount in situ hybridization for *Ntf3* in E12.5 control (*Left*) and *Sox2* cKO (*Right*) cochleae. (Scale bar, 100 µm.) *n* = 3 embryos per genotype.

**Fig. S4. Chromatin state and transcription-factor occupancy at the distal Bdnf elements D1 and D2.**

(A and B) Expanded views of D1 (A) and D2 (B); tracks as in Fig. 6 B and C. D1 carries H3K4me1 in both P1 hair cells and P1 supporting cells without H3K27ac in either, and overlaps neither a SOX2 footprint nor a SOX2 or ATOH1 peak. D2 is H3K27ac-marked in E12.0 prosensory cells and in P1 hair cells, retains H3K4me1 without H3K27ac in P1 supporting cells, and overlaps an E17.5 ATOH1 CUT&RUN peak. (Scale bars, 500 bp).

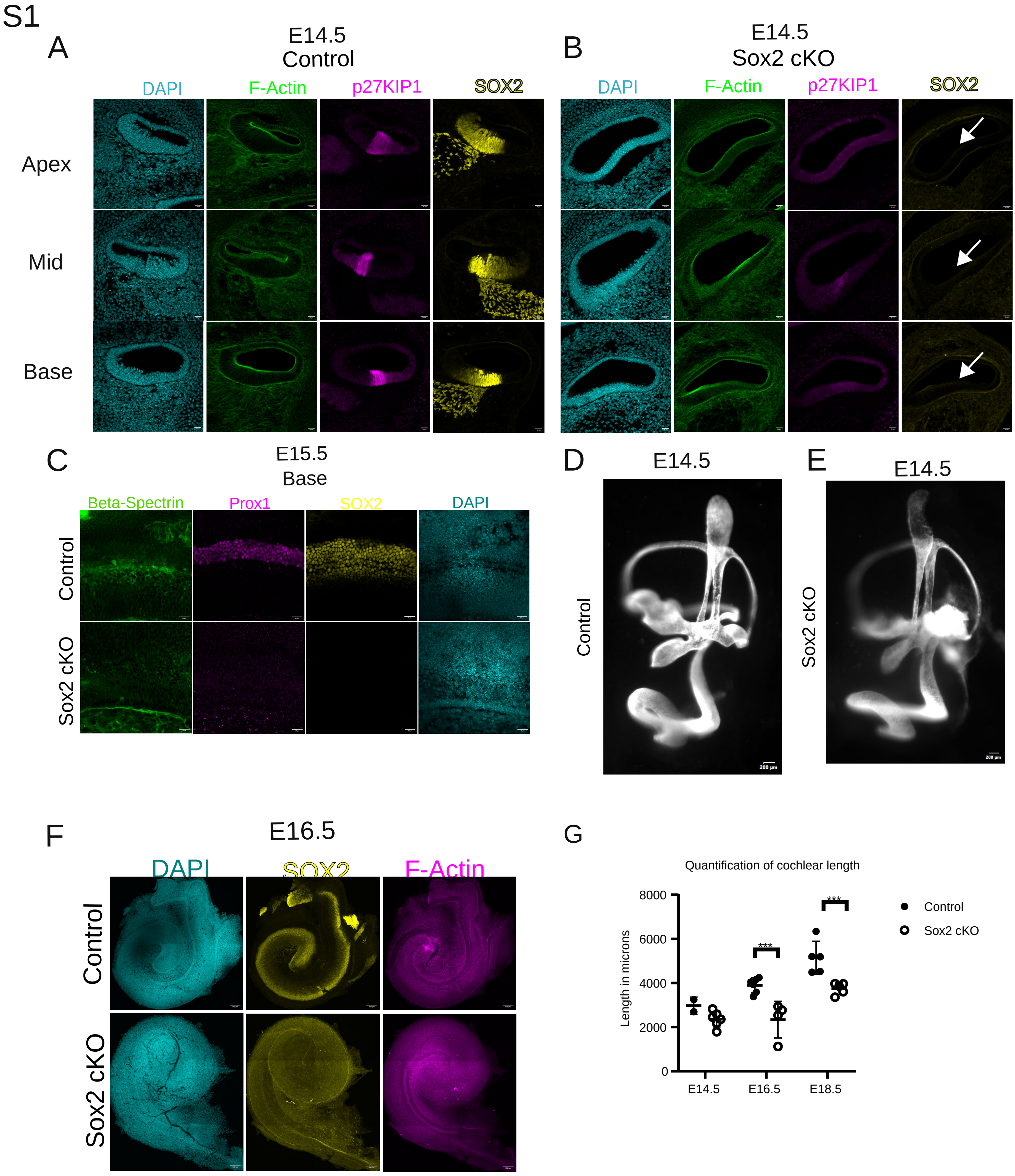
S1
A
E14.5
Control
DAPI
F-Actin
p27KIP1
SOX2
Apex
Mid
Base
B
E14.5
Sox2 cKO
DAPI
F-Actin
p27KIP1
SOX2
C
E15.5
Base
Beta-Spectrin
Prox1
SOX2
DAPI
Control
Sox2 cKO
D
E14.5
Control
200 µm
E
E14.5
Sox2 cKO
200 µm
F
E16.5
DAPI
SOX2
F-Actin
Control
Sox2 cKO
G
Quantification of cochlear length
Length in microns
8000
6000
4000
2000
0
E14.5
E16.5
E18.5
***
***
Control
Sox2 cKO

S2

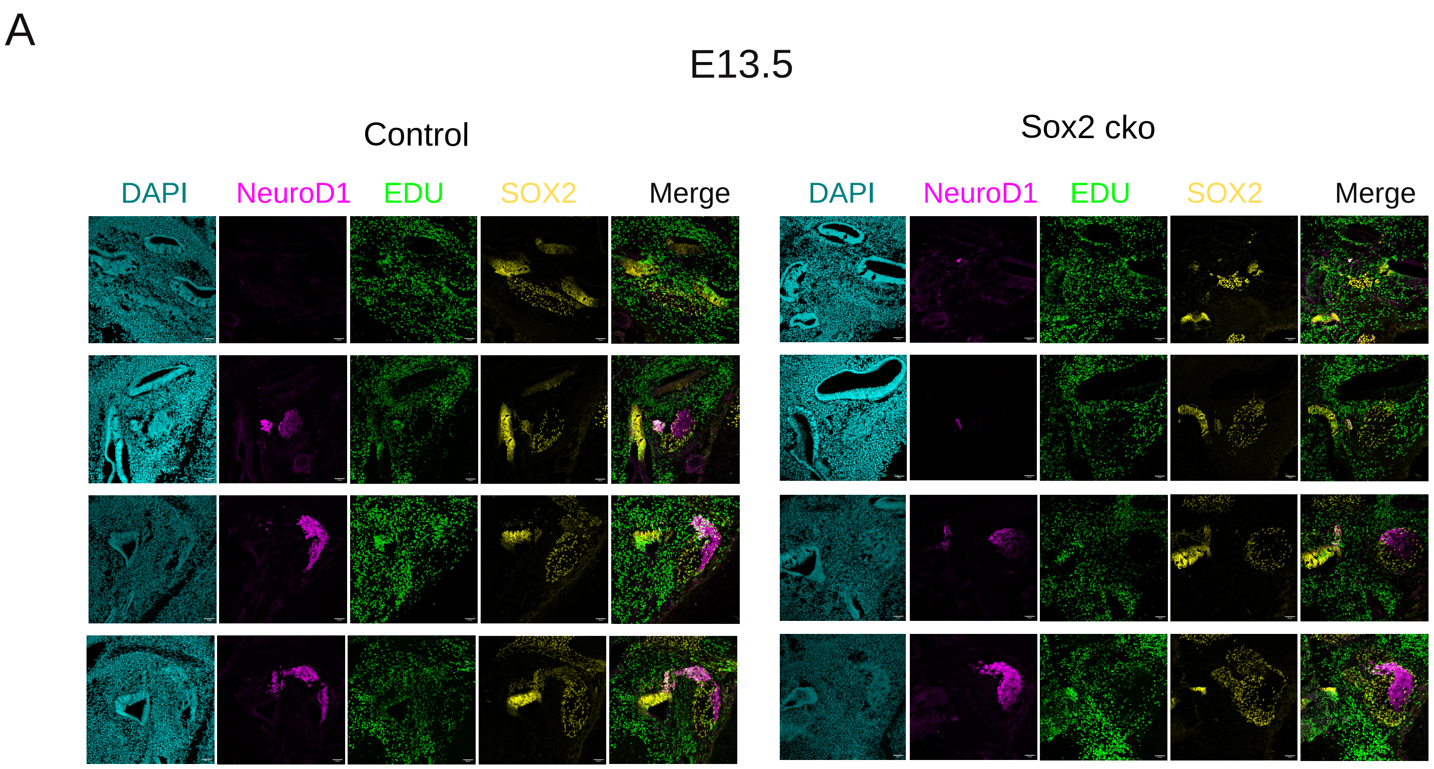
A
E13.5
Control
Sox2 cko
DAPI
NeuroD1
EDU
SOX2
Merge
DAPI
NeuroD1
EDU
SOX2
Merge

S3

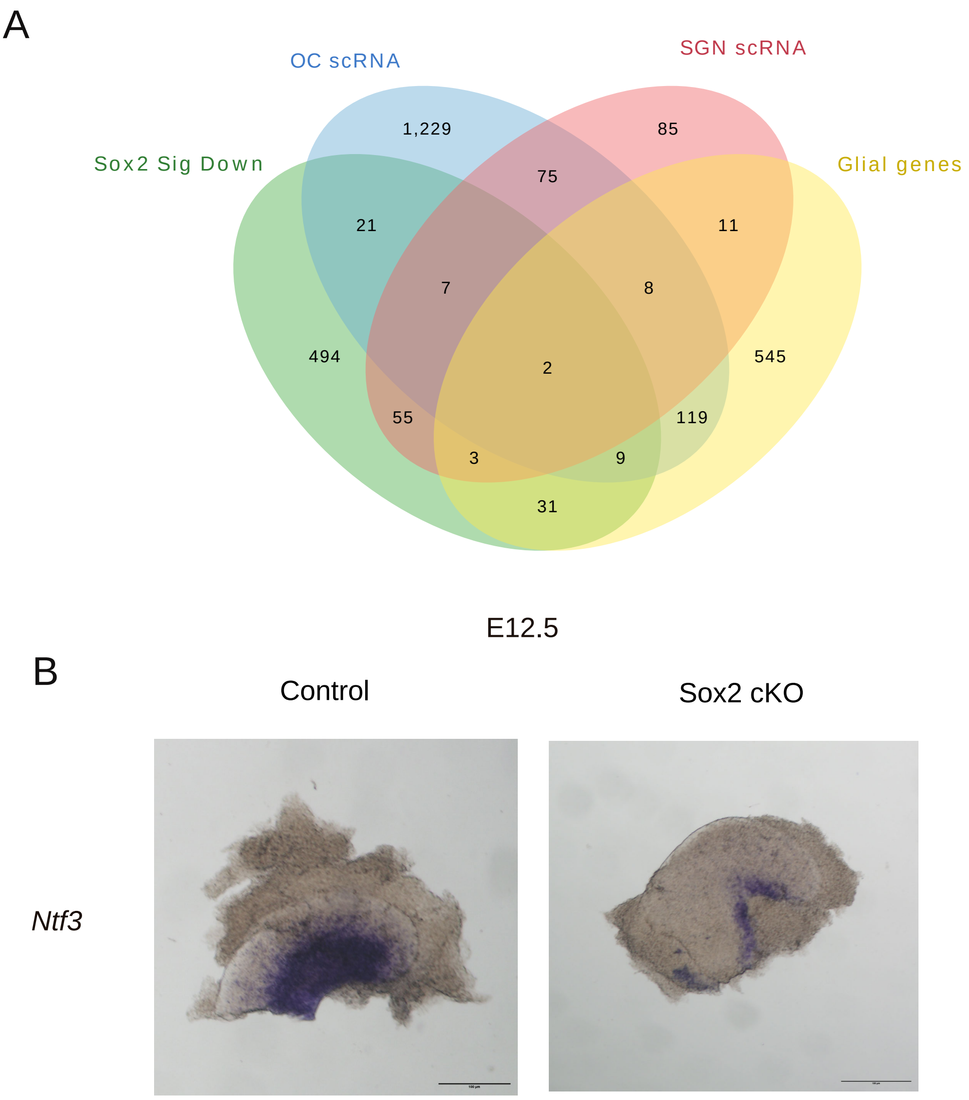
A
OC scRNA
SGN scRNA
Sox2 Sig Down
Glial genes
1,229
85
75
21
11
7
8
494
2
545
55
119
3
9
31
E12.5
B
Control
Sox2 cKO
Ntf3

S4

A

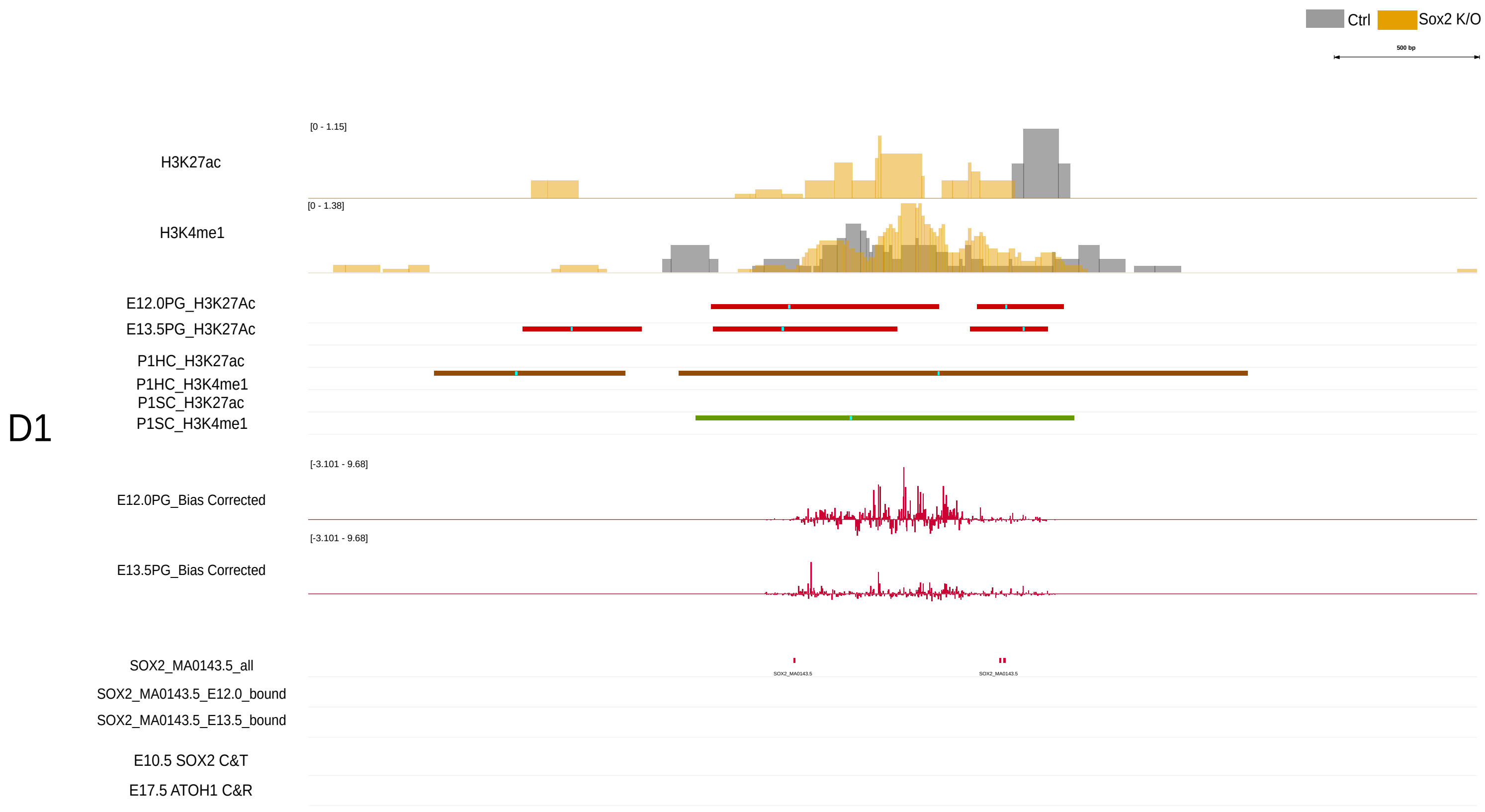
Ctrl
Sox2 K/O
500 bp
D1
H3K27ac
[0 - 1.15]
H3K4me1
[0 - 1.38]
E12.0PG_H3K27Ac
E13.5PG_H3K27Ac
P1HC_H3K27ac
P1HC_H3K4me1
P1SC_H3K27ac
P1SC_H3K4me1
E12.0PG_Bias Corrected
[-3.101 - 9.68]
E13.5PG_Bias Corrected
[-3.101 - 9.68]
SOX2_MA0143.5_all
SOX2_MA0143.5_E12.0_bound
SOX2_MA0143.5_E13.5_bound
E10.5 SOX2 C&T
E17.5 ATOH1 C&R

B

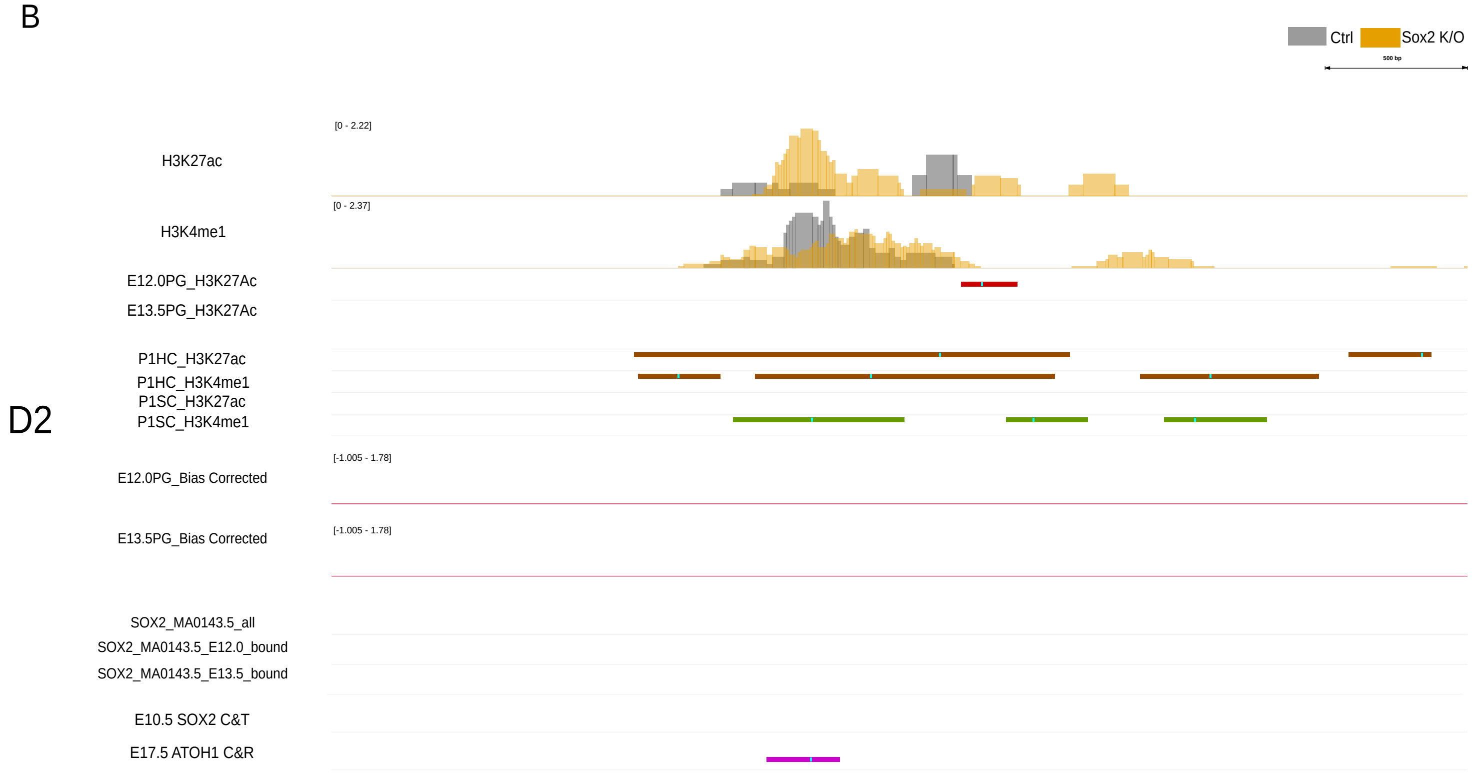
Ctrl
Sox2 K/O
500 bp
D2
H3K27ac
[0 - 2.22]
H3K4me1
[0 - 2.37]
E12.0PG_H3K27Ac
E13.5PG_H3K27Ac
P1HC_H3K27ac
P1HC_H3K4me1
P1SC_H3K27ac
P1SC_H3K4me1
E12.0PG_Bias Corrected
[-1.005 - 1.78]
E13.5PG_Bias Corrected
[-1.005 - 1.78]
SOX2_MA0143.5_all
SOX2_MA0143.5_E12.0_bound
SOX2_MA0143.5_E13.5_bound
E10.5 SOX2 C&T
E17.5 ATOH1 C&R
